# The rapid radiation of *Bomarea* (Alstroemeriaceae: Liliales), driven by the rise of the Andes

**DOI:** 10.1101/2022.09.15.507859

**Authors:** Carrie M. Tribble, Fernando Alzate-Guarín, Etelvina Gándara, Araz Vartoumian, J. Gordon Burleigh, Rosana Zenil-Ferguson, Chelsea D. Specht, Carl J. Rothfels

**Affiliations:** University Herbarium and Department of Integrative Biology University of California, Berkeley, CA 94709, USA; School of Life Sciences, University of Hawai’i at Mānoa, Honolulu, HI, 96822, USA; Grupo de Estudios Botánicos (GEOBOTA) and Herbario Universidad de Antioquia (HUA), Instituto de Biología, Facultad de Ciencias Exactas y Naturales, Universidad de Antioquia, Medellín, Colombia; Facultad de Ciencias Biológicas, Benemérita Universidad Autónoma de Puebla, Puebla, Puebla, 72570, Mexico; Department of Oral Biology, University of California, Los Angeles, CA 90095, USA; Department of Biology, University of Florida, Gainesville, FL 32611 USA; Department of Biology, University of Kentucky, Lexington KY 40506 USA; Section of Plant Biology and the L.H. Bailey Hortorium, School of Integrative Plant Science, Cornell University, Ithaca, NY 14853 USA; Intermountain Herbarium, Department of Biology, and Ecology Center, Utah State University, Logan, UT 84322 USA

**Keywords:** Andean uplift, divergence-time estimation, evolutionary radiation, hyb-seq, Liliales, recent rapid radiation

## Abstract

Complex geological events such as mountain uplift affect how, when, and where species originate and go extinct, but measuring those effects is a longstanding challenge. The Andes arose through a series of complex geological processes over the past c. 100 million years, impacting the evolution of regional biota by creating barriers to gene flow, opening up new habitats, and changing local climate patterns. *Bomarea* are tropical geophytes with ranges extending from central Mexico to central Chile. Of the roughly 120 species of *Bomarea*, most are found in the Andes, and previous work has suggested that *Bomarea* diversified rapidly and recently, corresponding with the uplift of the Andes. While many *Bomarea* species occur over small, isolated ranges, *Bomarea edulis* occurs significantly beyond the ranges of any other *Bomarea* species (from central Mexico to northern Argentina) and is thought to have potentially human-mediated dispersal, due to its status as a pre-Columbian food plant. To untangle the potential drivers of diversification and biogeographic history in *Bomarea*, we used a target-capture approach to sequence nuclear loci of 174 accessions of 124 species, including 16 outgroup species from across the family (Alstroemeriaceae). We included 43 individuals of *B. edulis* from across its range to assess species monophyly and identify infraspecific phylogeographic patterns. We model biogeographic range evolution in *Bomarea* and test if Andean orogeny has impacted its diversification. We find that *Bomarea* originated in the central Andes during the mid-Miocene, then spread north, following the trajectory of major mountain uplift events. Most observed speciation events occurred during the Pleistocene, while global climate cooled and oscillated and the northern Andes achieved their current form. Furthermore, we find that Andean lineages diversified faster than their non-Andean relatives. These results demonstrate a clear macroevolutionary signal of Andean orogeny on this neotropical radiation.

## Introduction

Beginning with the work of early naturalists, such as Alexander von Humboldt’s documentation of the turnover of plant communities over an elevational gradient in the Andes (Von Humboldt and Bonpland, 1807) and Alfred Russel Wallace’s observations on the connection between geographic barriers and species distributions (Wallace, 1863), evolutionary biologists have recognized the importance of geological processes on generating and maintaining biodiversity. Despite the central role of these processes in biodiversity dynamics, key methodological and empirical challenges remain in understanding how lineages respond to events such as continental drift, large-scale climatic changes, and mountain uplift. Tropical America—one of the most biodiverse regions on the planet—provides a key opportunity to study these interactions. Specifically, Andean uplift, which has created abiotic barriers and opened up novel habitats, is thought to be one of the main drivers of rapid radiations leading to high species-richness and endemism (Koscinski et al., 2008; Guayasamin et al., 2017; Ceccarelli et al., 2016; Ribas et al., 2007; Chazot et al., 2018; Lisa De-Silva et al., 2017; Madriñán et al., 2013; Lagomarsino et al., 2016; Hughes and Eastwood, 2006). However, tropical regions worldwide, including tropical America, are understudied when compared to temperate regions (Collen et al., 2008; Titley et al., 2017), and the evolutionary histories of many tropical American clades are unknown (Pérez-Escobar et al., 2022). Without detailed and reliable phylogenies, accurate inference of the effect of geological events on evolutionary processes is impossible. In this study, we built the first well-sampled phylogeny of the primarily Andean plant genus *Bomarea* Mirb. and modeled how this clade has spread across South America in the context of tectonic movements, climatic change, and Andean uplift.

The Andes is over 9000 kilometers long and its tallest mountain (Aconcagua in Argentina) reaches 6962 meters above sea level, second only to Asian mountains such as the Mount Everest in the Himalayas (Graham, 2009). The emergence of the Andes occurred over the past ∼100 million years (myr) through geologically complex uplift that modified continent-scale topography, climate, watersheds, habitats, and, correspondingly, biota (Graham, 2009; Hoorn et al., 2010; Pérez-Escobar et al., 2022). The tropical Andes is a global biodiversity hotspot with roughly 20,000 endemic plant species and 1,557 endemic vertebrate species (Myers et al., 2000). While evolutionary diversity peaks at mid elevations in the Andes (Griffiths et al., 2021), high-elevation habitats have high rates of endemism and lineages that are often marked by fast rates of evolution, perhaps due to the recent emergence of such habitats or to increased mutation rates caused by high UV exposure (Madriñán et al., 2013). Uplift began in the southern Andes and spread north; thus, the northern Andes are the youngest and most topographically complex part of the range, while the southern Andes reach higher elevations (Graham, 2009; Hoorn et al., 2010; Pérez-Escobar et al., 2022). Over this period, the rise of the Andes affected regional and global climate by creating a steep precipitation gradient from west (dry) to east (moist) across South America and by sequestering CO_2_ through increased erosion of exposed silicate rock, which contributed to global cooling during the late Cenozoic (Graham, 2009). Andean orogeny led to the creation and then drainage of a massive wetland (Pebas system) over much of the current extent of the Amazon basin (Hoorn et al., 2010). Mountainbuilding also opened up novel tropical alpine habitat, allowing for the emergence of new ecosystems like paramo and puna (Madriñán et al., 2013; Pérez-Escobar et al., 2022), and increased topographic complexity through the formation of mountain peaks and deep river valleys that shape spatial biodiversity patterns (Hazzi et al., 2018).

There are three primary patterns of spread and colonization in Andean clades; these patterns have different implications for how uplift may have affected the biodiversity dynamics of local biota. First, clades may have spread from north (North or Central America) to south, dispersing over the oceanic gap between Central and South America or via the Isthmus of Panama after its formation. Since the northern Andes is the youngest part of the Cordillera, north-to-south clades with strictly montane distributions likely arrived to the Andes once northern South America had already sustained significant mountain building activity. These clades may subsequently disperse south along the already-formed Andean chain. The contribution of northern-temperate flora to the current composition of high-elevation Andean ecosystems has been discussed by several previous studies (Sklenář et al., 2011; Simpson, 1975; Bacon et al., 2018). Notably, the plant genus *Lupinus* (Fabaceae) arrived to the Andes from North America and then radiated, demonstrating extremely high rates of speciation and morphological change over a very short time period (1.18–1.76 myr; Hughes and Eastwood, 2006). Additionally, *Gentianella* (Gentianaceae) is hypothesized to have dispersed from North America to northern South America after the emergence of alpine conditions and likely also the formation of the Isthmus of Panama (von Hagen and Kadereit, 2001). This pattern implies that the taxa did not evolve as the Andes formed; rather, they dispersed through and/or adapted to existing Andean habitats.

Second, clades may have originated in the south (southern South America) and spread northward, following the emergence of high-elevation habitats as the Andes formed. This pattern appears to be less common (at least in botanical systems) than north-to-south dispersal (Bacon et al., 2018). However south-to-north dispersal is evident in the iconic Andean *Puya* (Bromeliaceae), which originated in modern-day Chile and spread north, presumably as suitable habitat emerged during Andean uplift (Jabaily and Sytsma, 2013). *Chiquiraga* (Asteraceae) also originated in southern South America, subsequently spreading north (Ezcurra, 2002), and Lagomarsino et al. (2016) hypothesize a similar pattern underlies the centropogonid radiation in Andean cloud forests. Other examples include *Gunnera* (Bacon et al., 2018) and wax palms (*Ceroxylon*, Sanín et al., 2016). This pattern is reminiscent of the progression rule of island biogeography, in which clades first arrive to older islands and spread to newer ones as they are formed (as exemplified by many Hawaiian taxa: Hennig, 1966; Funk and Wagner, 1995). This pattern implies that the geological history of Andean uplift had measureable impacts on how, when, and where species formed.

Third, clades may have diversified in-situ, adapting to higher elevation habitats as they emerged and/or forming new species in response to increased topographic complexity and new dispersal barriers. High-elevation species also experience greater dispersal barriers, as their preferred habitats—high mountain peaks—are often enmeshed within a lower elevation matrix, creating an island-like system. For example, Ceccarelli et al. (2016) conclude that Andean uplift generated new habitats, which promoted adaptation and specialization to harsh, high-elevation habitats, and created dispersal barriers between high-elevation habitats, in *Brachistosternus*, a genus of scorpions. Andean uplift also affected the distributions and diversification of parrots (*Pionus*, Ribas et al., 2007). *Pionus* originated in lowland South America prior to Andean uplift. The Andes then split the clade into three clades: distinct highland (in the Andes), dry lowland (west of the Andes), and wet lowland (east of the Andes) lineages. Subsequently, climatic oscillations in the Pleistocene invigorated speciation in the highland parrots.

Distinguishing between these patterns requires accurate phylogenies on which to base comparative analyses, and yet recent rapid radiations introduce complex challenges for phylogenetic inference (Giarla and Esselstyn, 2015). Specifically, recent rapid radiations are characterized by rapid consecutive phylogenetic divergences—and thus very short internal branches. These branches are difficult to resolve due to the combined effects of low signal (there is little time for substitutions to accumulate) and rampant incomplete lineage sorting (ILS; Maddison, 1997). These challenges are exacerbated when working with non-model organisms with limited genomic resources, as it may not be possible to generate information about the variability of gene regions across individuals/species, or the presence of recent gene duplications that could affect homology inference, prior to sequencing.

In this study, we illustrate solutions to these challenges using *Bomarea*—a recent and rapid Andean radiation— target-capture approaches to data generation, and an analytical pipeline focused on the homology issues inherent in recent rapid plant radiations. We estimate the first well-sampled phylogeny of *Bomarea* and demonstrate how to recover and evaluate informative loci from universal probe sets, despite high rates of gene duplication and rapid recent divergences. We then infer divergence times and model the biogeographic history and diversification rates of the clade in the context of climatic and geological changes in South America over the past 80 million years. Our study thus adds to the current understanding of how the shifting tropical and southern American landscape contributed to the extremely high rates of extant species richness.

## Bomarea

*Bomarea* is one of four genera in Alstroemeriaceae (Liliales), and comprises roughly 120 described species, most of which occur in cloud forests and tropical alpine habitats such as paramo and puna regions throughout the Andean cordillera (Hofreiter, 2005; Alzate et al., 2008b). While many *Bomarea* are perennial twining vines, some species—mostly those that occur in alpine habitats above the tree line—are erect or suberect perennial herbs (Fig. 1; Alzate Guarín, 2005). Each above-ground stem senesces after flowering, and below-ground stems (rhizomes) and tuberous roots fuel the regrowth of subsequent above-ground structures (Tribble et al., 2021a,b).

**Figure 1:**
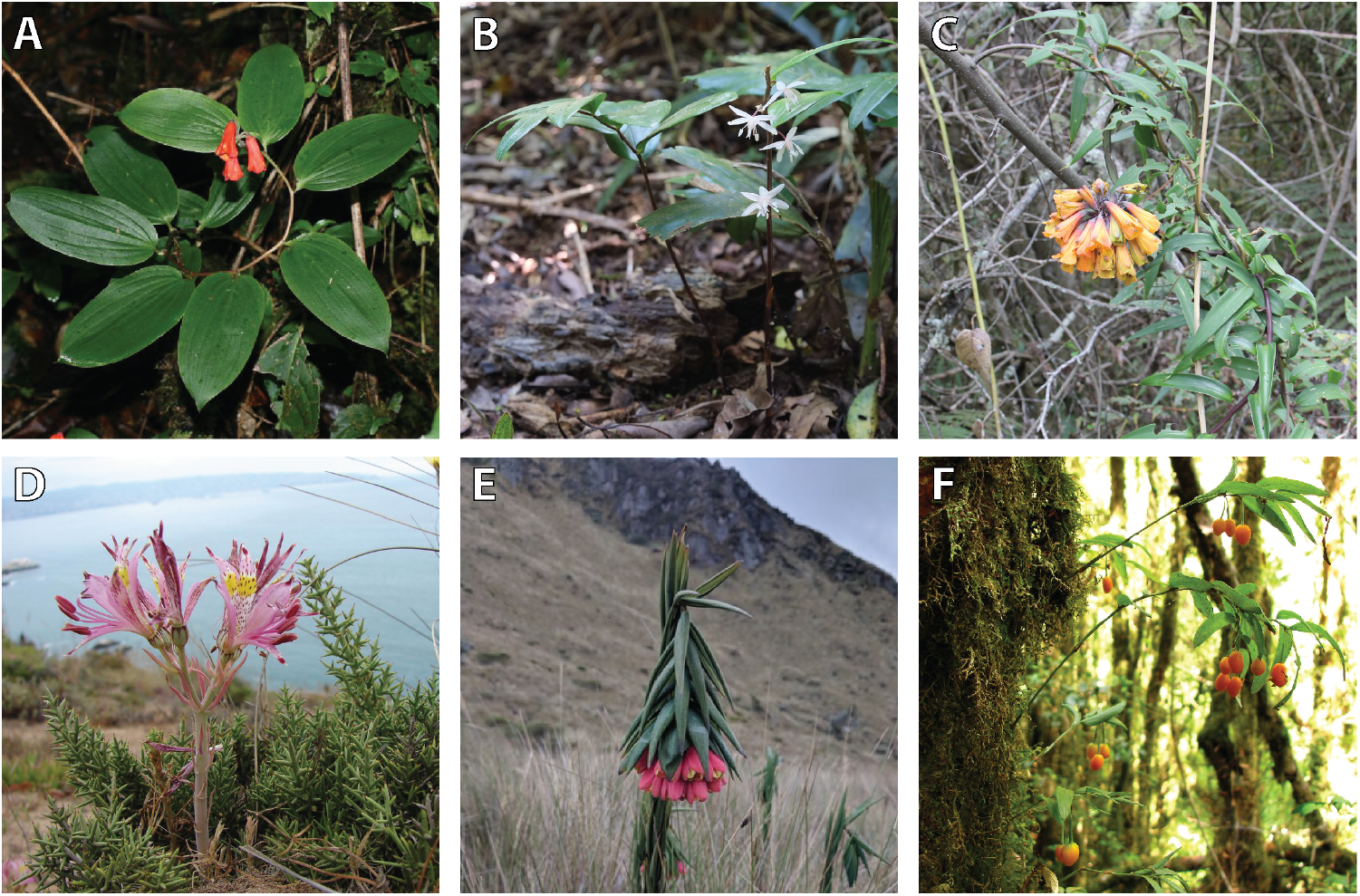
Species in Alstroemeriaceae: (A) *Bomarea suberecta*, photo: Robbin Moran; (B) *Drymophila moorei*, photo: Igor Makunin; (C) *Bomarea multiflora*; (D) *Alstroemeria hookeri*, photo: Patricio Novoa Quezada; (E) *Bomarea glaucescens*, photo: iNaturalist user fishy21, and; (F) *Luzuriaga radicans*, photo: Inao Vasquez.

Alstroemeriaceae likely originated in the late Cretaceous when Australasia (Australia and New Zealand), Antarctica, and southern South America were very close or still physically connected (Chacón et al., 2012). Of the four genera in Alstroemeriaceae, *Bomarea* and sister genus *Alstroemeria* are restricted to the Americas while two other genera occur in Australasia: *Luzuriaga* occurs in southern South America and New Zealand, and *Drymophila* is restricted to Australia. *Bomarea* occurs from Chile and Argentina to Mexico and the Caribbean (Alzate et al., 2008a). While many species have highly restricted ranges, *Bomarea edulis* (Tussac) Herb. is found throughout the entire range of *Bomarea*. Furthermore, *B. edulis* is the only species of *Bomarea* in the Caribbean, the northern-most occurring species in central Mexico, the only species in Brazil, and its populations in northern Argentina extend the range of the genus to the south. Substantial morphological variability across its range led to the recognition of at least 23 species within *B. edulis*. However, current taxonomies reduce these all to synonymy under B. edulis (Hofreiter, 2006). This species produces edible tubers and was previously cultivated (Hofreiter, 2006), leading to the hypothesis that its wide distribution might be due to human-mediated dispersal. Despite this taxonomic and biogeographic confusion, no previous study has gathered molecular data to examine relationships within *B. edulis*, including determining if *B. edulis* is monophyletic. If “*B. edulis*” as currently circumscribed does indeed encompass several morphologically similar species, then the geographic range of those species and their placement within the *Bomarea* phylogeny could greatly influence biogeographic inference within the genus. Conversely, if *B. edulis* is in fact a single monophyletic species, then placing that species within the *Bomarea* phylogeny will facilitate modeling its range expansion in the context of biogeographic movements across the group.

Our current understanding of the *Bomarea* phylogeny derives largely from Alzate et al. (2008a) and from Chacón et al. (2012), which also addressed biogeographic patterns in *Bomarea*. However, Chacón et al. (2012) did not reconstruct the finer-scale movements of species in *Bomarea*, and relationships within the genus—including of the single sampled *B. edulis* individual—were highly uncertain. This was likely due to the lack of variable markers, which has also limited other previous attempts to reconstruct the genus’ phylogeny (*e*.*g*., Alzate et al., 2008a). Several previous studies have hypothesized that *Bomarea* evolved in the context of recent Andean uplift because of the tendency of species to thrive in montane regions and because the center of species diversity is in the central Andes (Alzate et al., 2008a; Hofreiter, 2007). Additionally, Chacón et al. (2012) found that *Bomarea* diversified beginning 14.3 million years ago (Ma) during a period of ongoing Andean orogeny. However, sparse taxon sampling and limited taxonomic resolution prevented the authors from addressing biogeographic dynamics in the genus.

## Materials and Methods

### Sample Collection and Sequencing

We obtained samples from 192 individuals from silica-dried and herbarium-sampled material (Table S1). These samples represented 174 *Bomarea*, 14 *Alstroemeria*, two *Drymophila*, and three *Luzuriaga* individuals comprising 137 different species. We extracted DNA from all samples using a standard CTAB protocol (Doyle, 1991). For the target enrichment, we used the GoFlag angiosperm 408 probes, which cover 408 relatively conserved exons that are found in a total of 226 single or low copy nuclear genes Endara and Burleigh (2022). The library preparation, target enrichment, and sequencing were all done by RAPiD Genomics (Gainesville, FL) using protocols described in Breinholt et al. (2021) and Endara and Burleigh (2022) (see Supplemental Section 1.1 for details). Samples were sequenced on an Illumina HiSeq 3000 with 100 base-pair (bp) paired-end reads.

### Data Processing

We assembled multiple sequence alignments from the target regions using the bioinformatic pipeline described in (Endara and Burleigh, 2022), which modified the pipeline from Breinholt et al. (2021) for angiosperms. This pipeline conducts a de novo assembly with BRIDGER version 2014-12-01 (Chang et al., 2015) based on sequence homology of raw reads to a set of reference sequences for each locus. We aligned the recovered sequences from the target regions only from each locus using `MAFFT` version 7.425 (Katoh and Standley, 2013) and merged any isoforms from the same taxon with heterozygous sites a Perl script that used IUPAC ambiguity codes to represent sites with multiple nucleotides. This pipeline does not explicitly phase loci; however, in some cases, a locus alignment might include multiple sequences from some samples, representing cases in which the BRIDGER assembler identified what is likely more than allelic diversity. We discarded all by-catch and aligned only targeted regions, and not flanking sequences, to minimize missing data and misaligned regions. For all loci, we retained all recovered copies per individual.

We then further processed the alignments to remove possible contaminants, minimize missing data, and address homology issues within loci. We performed all downstream processing in R (v4.1.0, R Core Team, 2013) unless otherwise noted. We first discarded loci that were recovered from fewer than a third (57) of all individuals. We then built gene trees of the remaining loci with `IQtree` (Nguyen et al., 2015) using `ModelFinder` (Kalyaanamoorthy et al., 2017) to identify the best nucleotide model. We manually inspected all gene trees to check for possible contaminant sequences—those that were on long branches or from ingroup individuals that fell with the outgroup. We then blasted (blastn, Altschul et al., 1990) each potential contaminant sequence and removed those whose best blast hit was not another monocot. We then re-built all gene trees using the same method as above.

For those gene trees retaining some multiple copies per individual for at least some loci after the “decontamination”, above, we manually inspected each tree and attempted to resolve each case of multiple copies per individual. We first determined if the duplicated loci appeared due to putative gene duplication across all individuals—, gene duplication of only a few individuals, or impossible to determine (see Fig. 2). For gene duplication events across all or most individuals, we split the tree into multiple reciprocally monophyletic clades (Fig. 2 A). For gene duplication of only a few individuals, we retained no more than one sequence per individual by removing one sequence randomly if tips were sister (Fig. 2 B) or removing both if placement was uncertain. For any indeterminate case we removed the locus entirely. We then reflected all changes to the gene trees onto the original alignments; we split alignments if we had identified multiple orthologous clades in the gene trees and we removed all sequences from alignments that we had removed from the gene trees. Finally, we removed any individual from an alignment if it had less than 10% matrix occupancy (more than 90% missing data). All downstream analyses were performed on these cleaned alignments.

**Figure 2:**
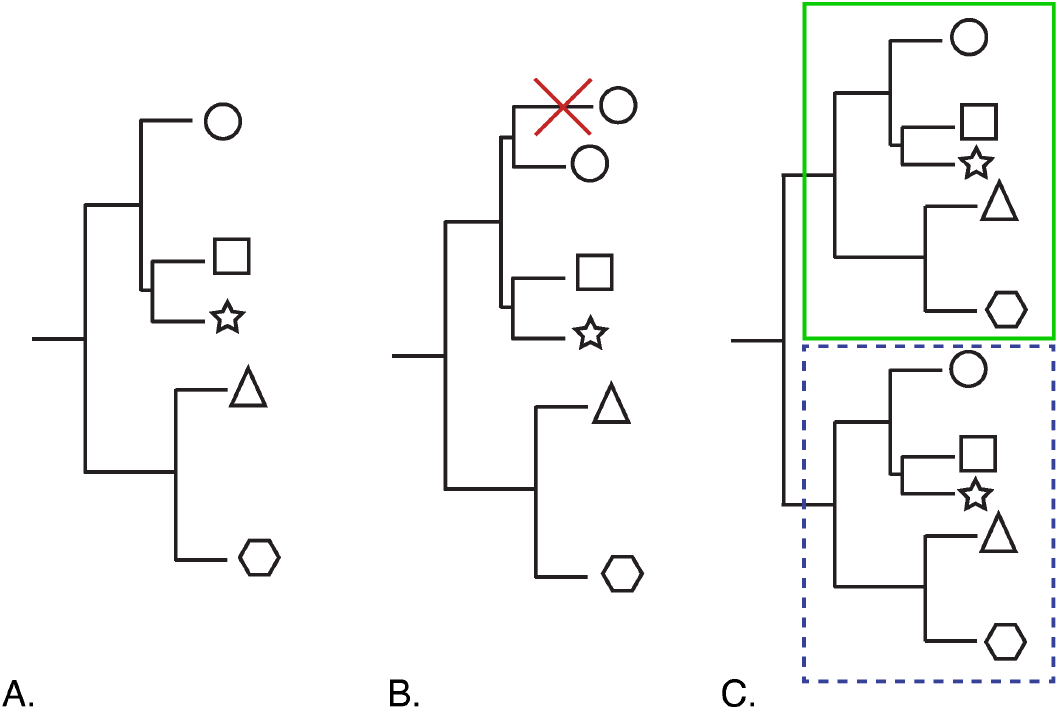
Possible patterns of putative gene duplication (hereafter, gene duplication) in loci. Shapes at tips represent individuals. (A) No gene duplicates in the locus—all tips represent unique individuals). (B) Local gene duplication—duplicates monophyletic. (C) Ancestral gene duplication—locus can easily be split into two clades within which tips represent unique individuals.

### Phylogenetic Reconstruction

We inferred phylogenetic relationships using two methods: a “species tree” methods based on an approximate multispecies coalescent approach (A-MSC)—`ASTRAL-II` (Zhang et al., 2018)—and one maximum likelihood concatenate supermatrix approach implemented in `IQtree` (Nguyen et al., 2015).

For the A-MSC analysis, we built gene trees of each locus, again using `IQtree` with 100 standard bootstrap replicates, and using `ModelFast` to infer the best nucleotide model for each locus. We collapsed all nodes with *<* 10% bootstrap support in each gene tree and ran `ASTRAL-II` with default settings, treating each individual as its own species (*i*.*e*., as a tip in the resulting phylogeny).

For the ML analysis, we concatenated all loci and used `PartitionFinder 2` (Lanfear et al., 2017) to determine the best configuration of partitions given the option to partition by locus and by codon position and the best nucleotide evolution model(s) for those partitions. We used AICc (Bedrick and Tsai, 1994) for model selection and performed a greedy search. We used the inferred partitioning scheme and models for ML analysis with IQtree, implemented on the CIPRES computing platform (Miller et al., 2011). We also ran 1,000 ultrafast bootstrap replicates (Hoang et al., 2018).

Given the short divergences between many nodes in the trees, ILS may be an important process, and thus we used the A-MSC topology from the `ASTRAL-II` analysis for all subsequent analyses.

### Divergence-Time Estimation

We dated the A-MSC topology in Revbayes (Höhna et al., 2014) using an exponential relaxed clock model, birthdeath tree prior, and a partitioned *GTR* + Γmodel of nucleotide substitution. Running the divergence-time estimation (DTE) on the full set of loci was not computationally feasible, so we used the `R script genesortR` (Mongiardino Koch, 2021) to select a set of 15 loci; we partitioned these data by locus and codon position for a total of 45 subsets. In genesortR we defined the ingroup as *Alstroemeria* plus *Bomarea*, only considered loci with *>*10% of ingroup terminals, removed outliers (1% of loci that differ most in PCA-space), and set `topological similarity` to “true” (see hrefhttps://github.com/mongiardino/genesortR for more details on these parameters). `genesortR` sorts remaining loci by a PCA axis of “phylogenetic usefulness” that takes into account potential biases (*e*.*g*., average pairwise patristic distance) and potential beneficial qualities (*e*.*g*., average bootstrap support); we then selected the top 15 loci.

Our birth-death prior on the tree assumes tips represent species—the result of a dichotomous speciation and extinction process. To accommodate this assumption, if a species was represented by a multiple tips forming a clade, we selected one tip at random and removed the others. When tips identified as the same species were not monophyletic, we kept one random representative of each clade.

Because the `RevBayes` DTE analysis requires a starting tree that has a non-zero probability given the node calibrations (Table 1), we first dated the A-MSC topology using penalized likelihood in R with the chronos() function from the ape package (R Core Team, 2013; Paradis and Schliep, 2019). We then used this chronogram as the starting tree in our subsequent relaxed-clock analysis in `RevBayes`.

**Table 1:**
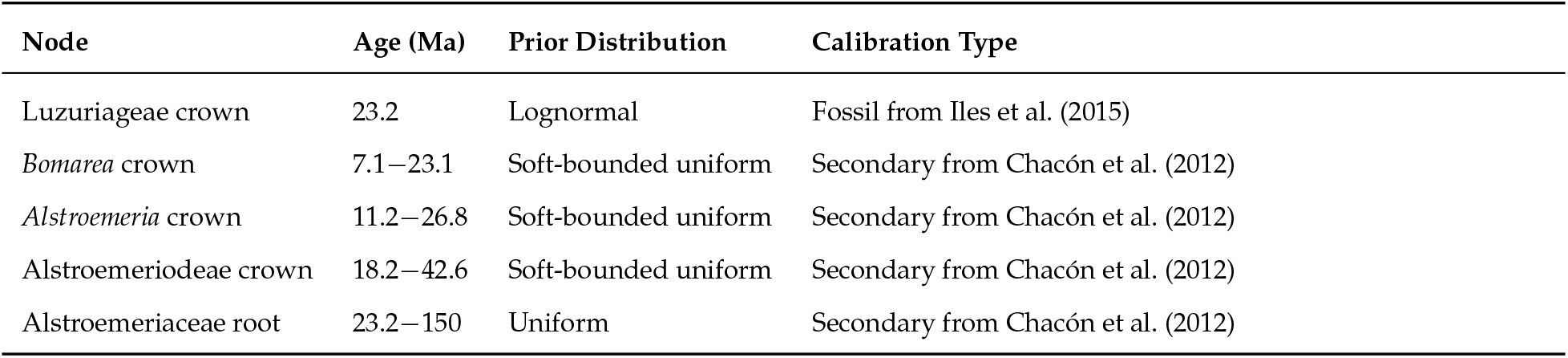
Calibrations and constraints used for divergence-time estimation. Normal soft-bound uniform distribution refers to a prior distribution on parameter values, where each “end” of the uniform distribution ends with a half-normal distribution rather than an abrupt change in probability (similar to Yang and Rannala, 2006). More details are available in `RevBayes` code on GitHub.

| Node | Age (Ma) | Prior Distribution | Calibration Type |
| --- | --- | --- | --- |
| Luzuriageae crown | 23.2 | Lognormal | Fossil from Iles et al. (2015) |
| *Bomarea* crown | 7.1–23.1 | Soft-bounded uniform | Secondary from Chacón et al. (2012) |
| *Alstroemeria* crown | 11.2–26.8 | Soft-bounded uniform | Secondary from Chacón et al. (2012) |
| Alstroemerioideae crown | 18.2–42.6 | Soft-bounded uniform | Secondary from Chacón et al. (2012) |
| Alstroemeriaceae root | 23.2–150 | Uniform | Secondary from Chacón et al. (2012) |

We calibrated the tree using one fossil calibration and three secondary calibrations from the most recent family-level analysis (Table 1, also see Chacón et al., 2012). We checked for convergence of estimated branch lengths and ages using Gelman and Rubin’s convergence diagnostic as implemented in `R` with the `gelman.diag()` function in the CODA package (Gelman and Rubin, 1992; Plummer et al., 2006).

To investigate the relative influence of molecular and fossil data on estimated node ages, we also ran a series of analyses. First, we ran the analysis under the “tree prior” with no calibrations (fossil or secondary) or molecular data, only our prior specifications on the tree (including a prior on the root age). We then ran the analysis without any molecular data but including the age of the fossil and secondary node calibration. Including these node calibrations represents a “calibration density”, which represents the amount of information about ages contained in the calibration densities alone.

### Biogeographic Range Reconstruction

We estimated biogeographic ranges at two taxonomic scales. First, we reconstructed the history across the full family-level dataset including the samples in Luzuriageae and *Alstroemeria*. To examine finer-scale patterns within *Bomarea* we also performed biogeographic reconstruction at on a *Bomarea*-only tree. We performed both analyses in `RevBayes` using a Dispersal-Extinction-Cladogenesis model of range evolution (Ree and Smith, 2008) over the maximum clade credibility (MCC) tree from the DTE as well as over a sample of 100 trees from the DTE posterior.

#### Alstroemeriaceae

We coded biogeographic ranges based on distribution data from the World Checklist of Selected Plant Families (WCSP, 2020). We then recoded species into five broad biogeographic regions: Australasia (Australia and New Zealand), southern South America (Chile, Argentina, and Uruguay), eastern South America (Guyana Shield and Brazil), the northern/ central Andean region (Bolivia, Peru, Ecuador, Colombia, and Venezuela, hereafter “Andean”), and Central America including Mexico and the Caribbean. We allowed a lineage to occupy no more than three areas simultaneously and only allowed dispersal into physically adjacent areas. Dispersal into and out of Australasia was allowed only through southern South America, as those regions were physically connected through Antarctica during the Cretaceous, and thus represents the most likely mode of transition between these currently-disconnected regions. We specified one dispersal rate and one extirpation rate irregardless of area.

#### Bomarea

As no *Bomarea* occur outside of the Americas, we did not include Australasia in the *Bomarea*-specific analysis. Instead, we distinguished between northern (Ecuador, Colombia, and Venezuela) and central (Peru and Bolivia) Andean regions. Similar to the full Alstroemeriaceae analysis, we also included Central America, eastern South America, and southern South America regions. We allowed lineages to occupy up to five areas simultaneously. As above, we only allowed dispersal through physically connected areas and we estimated one rate of dispersal and extirpation.

### Diversification-Rate Estimation

We estimated branch-specific net-diversification rates across *Bomarea* using a birth-death model that allows speciation and extinction rates to shift across the tree (Höhna et al., 2019). Our sampling of species within Alstroemeriaceae is heterogeneous across the family; the ratio of sampled species to known total species is much lower for *Alstroemeria* than for the other genera. To ensure that this discrepancy did not affect our rate estimates, we ran the analysis only on the *Bomarea* subtree (same phylogeny as used in the *Bomarea*-only biogeographic model).

We summarized the average diversification rate in Andean vs. non-Andean nodes. For each generation of the biogeographic *Bomarea*-only model, we classified the nodes as either Andean (end-state either central Andean, northern Andean, or both central and northern Andean) or non-Andean. We then selected a random generation from the branch-specific diversification rate analysis and recorded the sampled speciation and extinction rates for Andean and non-Andean nodes. We then calculated the average net-diversification rate (*speciation* − *extinction*) as the mean of sampled rates for the Andean nodes vs. non-Andean nodes and compute the difference. Once we had performed this calculation for all biogeographic model generations, we calculated the difference between Andean and non-Andean rates per generation. This difference, *d* represents the degree to which Andean nodes diversity faster, averaged over the uncertainty in biogeographic ancestral state estimates, assignment of rate categories to nodes, and speciation and extinction values of the rate categories.

We used Bayes factors to quantify the support for faster diversification rates in Andean lineages. The Bayes factor is computed as:

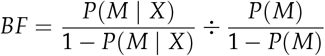

where *P*(*M*) is the prior probability that Andean lineages diversify faster, and *P*(*M* | *X*) is the posterior probability that Andean lineages diversify faster. We compute the posterior probability (*P*(*M* | *X*)) as:

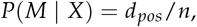

where *d*_*pos*_ is the number of differences that are greater than zero (*d >* 0) and *n* is the total number of differences (number of samples). Further, we assume the prior probabilities that Andean lineages diversify faster or slower are equal, *i*.*e*., *P*(*M*) = 1 − *P*(*M*), so that the Bayes factor is:

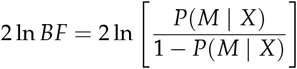

Positive values of 2 ln *BF* indicate support for faster Andean diversification.

### Plotting and R packages

We used `RevGadgets` (Tribble et al., 2022), `ggtree` (Yu et al., 2017), `ggplot2` (Wickham, 2011), and maps (Deckmyn et al., 2021) to plot results in `R`. We edited some figures with Adobe Illustrator.

## Data and Code Availability

All code is available on GitHub at https://github.com/cmt2/bom_phy_analysis. All data is available on Dryad at [XXXXX] and NCBI SRA [XXXXXX].

## Results

### Data Processing

Of the 192 individuals we extracted and submitted for sequencing, we obtained sequence data for some of the target loci from 172. Among these samples, the number of loci recovered ranged from one to 371, with an average of 146.6, and a median of 123. Across all samples, we recovered some sequence data from 403 of the 408 loci, 228 of which had sequence data from more than a third of the accessions. Of these 228, 56 had some accessions with multiple copies in that locus. After gene-tree inspection (Fig. 2), we removed 11 of those 56 from the analysis. We kept 30 after removing duplicate accessions (Fig. 2B), and we split 15 into multiple alignments (Fig. 2C).

We removed sequences from 161 loci of 54 accessions due to contamination: a total of 301 sequences. These sequences blasted to a variety of non-monocot taxa: 14 Enterobacterales, 1 Rhodospirillales, 12 fungi, 249 non-monocot plants, and 25 with no BLAST hits. The 5 most common contaminant genera were *Solanum* (tomato and relatives, 87 sequences), *Rosa* (rose and relatives, 55 sequences), *Fragraria* (strawberry and relatives, 22 sequences), and *Vitis* (grape and relatives, 7 sequences). We removed three accessions entirely, as they appeared contaminated in most gene trees (*B. macusani, B. huanuco*, and *B. schlerophylla*).

After these data cleaning steps, the final dataset included 121 accessions of 221 loci; when concatenated, the matrix contains 43,218 base pairs with 33.5% gaps.

### Phylogeny of *Bomarea*

We find strong support for monophyletic genera and for the currently accepted genus-level relationships in both AMSC and ML trees: *Bomarea* and *Alstroemeria* are sister to each other (Alstroemeriodeae), *Luzuriaga* and *Drymophila* are sister to each other (Luzuriageae), and the two subfamilies are also sister (Fig. 3). Within *Bomarea*, we infer a backbone topology where *Bomarea salsilla* Vell., *B. ovallei* (Phil.) Ravenna, *B. obovata* Herb., and *B. edulis* form a grade sister to the rest of *Bomarea* (core *Bomarea*). However, the approximate coalescent method (`ASTRAL-II`—Fig. S1 vs. the ML concatenated tree (Fig. S2) differ in the inferred order of the grade: in the approximate coalescent result *B. edulis* is sister to core *Bomarea* while in the ML results *B. obovata* is sister to core *Bomarea* (summarized in Fig. 3A). This major topological difference may indicate that ILS is prevalent in these data (see Discussion).

**Figure 3:**
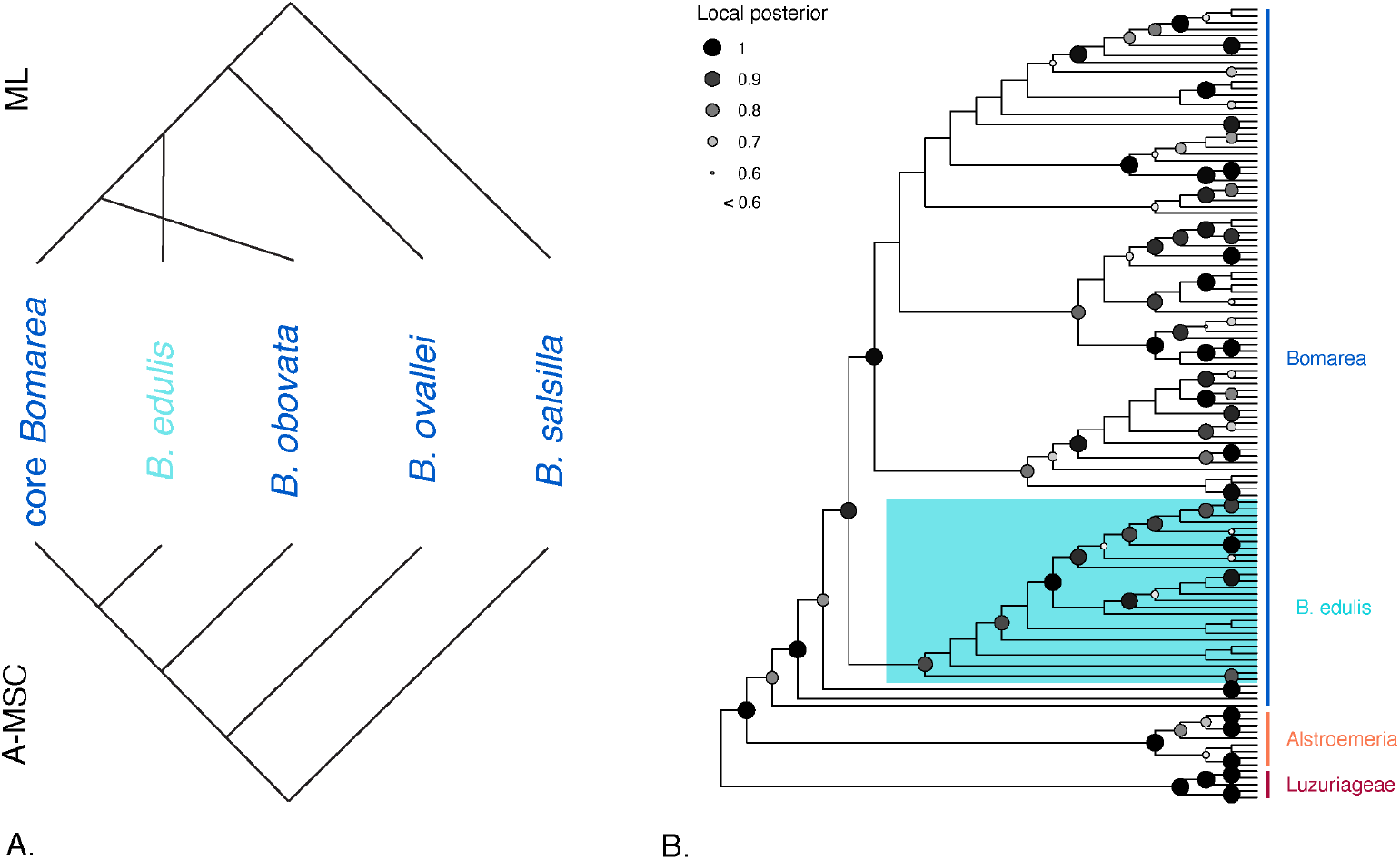
Topologies from species tree inference. (A) Topologies produced by Maximum Likelihood (ML) inference (concatenation) in IQtree (top) and by approximate multispecies coalescent (A-MSC) approaches in `ASTRAL-II` and `SVDq`. (B) Cladogram of Alstroemeriaceae showing broad relationships as inferred by `ASTRAL-II`.

We find strong support for a clade of most *B. edulis* individuals (Fig. 3, also true in the ML inferences—Fig. S2), though some individuals identified as *B. edulis* fall elsewhere (see Discussion). Despite strong support for the major relationships in the phylogeny, support remains low in several parts of the tree, in particular within core *Bomarea* at mid-aged nodes (local posteriors = 0.3–0.5, Fig. 3) and for many relationships within *B. edulis*. Because of a well-supported (*pp* = 0.88) divergence between two Argentinian individuals of *B. edulis* and the rest of the species’ clade (Fig. S1), we suspect that the Argentinian population may represent a separate species. We thus include two *B. edulis* individuals in the phylogeny for downstream DTE, biogeographic, and diversification rate estimation analyses: one individual from Argentina and one from Mexico.

### Divergence-time Estimation

Our DTE analysis infers a posterior mean estimation for the divergence of *Bomarea* and *Alstroemeria* (node 1 in Fig. 4) at 73.47 million years ago (Ma), during the Late Cretaceous. However, the 95% highest posterior density (HPD) interval for this node is broad (113.23–32.53 Ma), indicating considerable uncertainty in this estimate. Subsequently, *Luzuriaga* and *Drymophila* (node 2 in Fig. 4) diverged at 28.40 Ma (23.2–41.44) in the Oligocene and *Alstroemeria* and *Bomarea* (node 3 in Fig. 4) diverged at 21.76 Ma (15.34–28.89) during the early Miocene.

**Figure 4:**
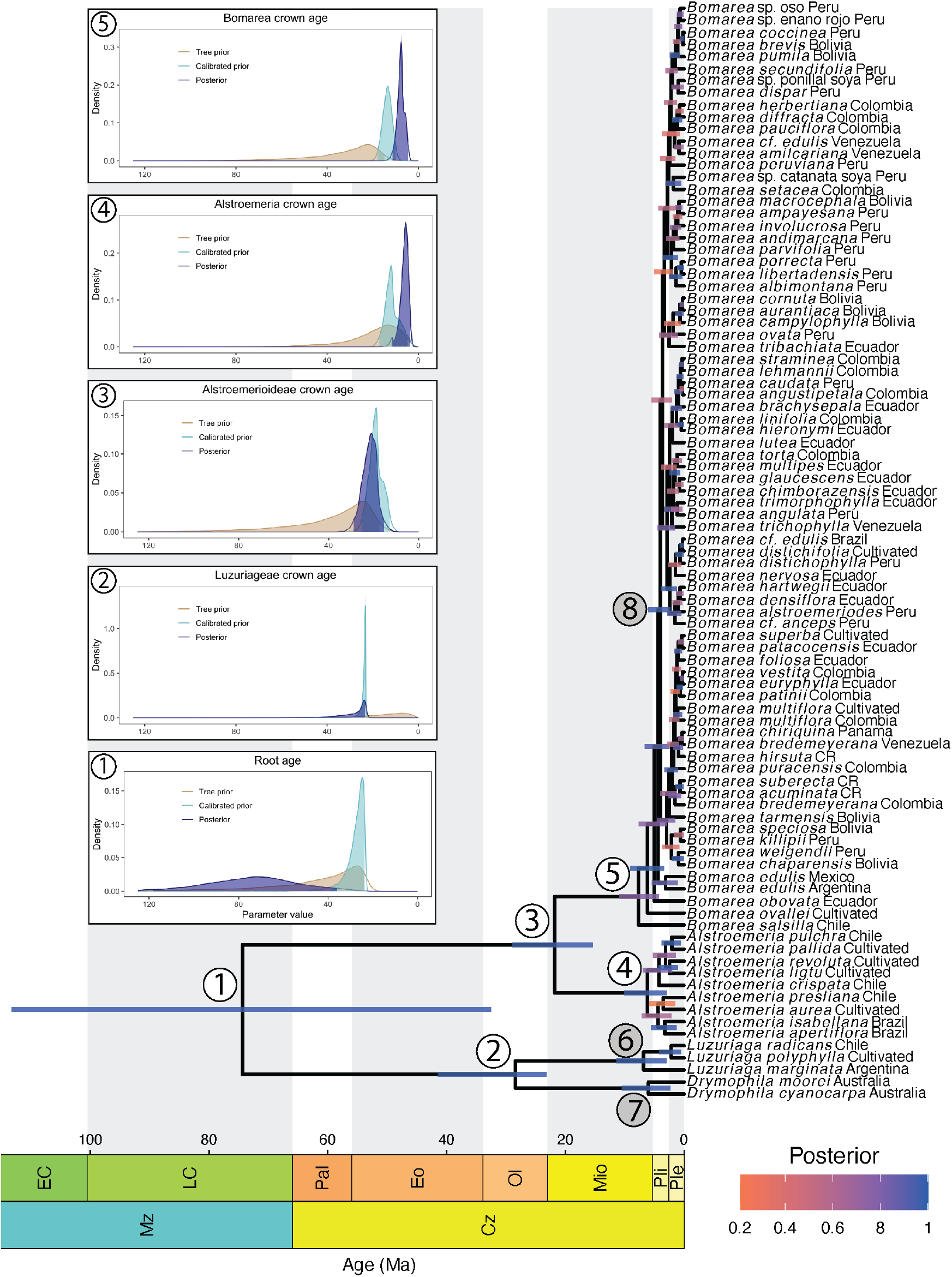
Chronogram of Alstroemeriaceae. Geological timescale shows age in millions of years ago (Ma). Node bars represent the 95% HPD for node ages. Color of the node bars indicates local posterior probability of nodes in the phylogeny (from `ASTRAL-II` analysis). Red asterisks indicate the four calibrated nodes. The four inset charts correspond to these calibrated nodes. In the insets, three probability distributions correspond to the ages estimated from the tree prior, calibrated prior (when the calibration ages are incorporated), and the posterior (when molecular data and calibration ages is incorporated).

We date the crown age of *Alstroemeria* (node 4 in Fig. 4) at 6.20 Ma (2.91–10.09 Ma) and the crown age of *Bomarea* (node 5 in Fig. 4) at 7.69 Ma (4.26–10.87 Ma), both in the Miocene, though our estimate of the *Alstroemeria* crown age may be biased (towards younger values) by the discrepancy between our highly-sampled *Bomarea* clade and less well-sampled *Alstroemeria* clade—it is possible that sample does not capture the crown node. Core *Bomarea* (node 8 in Fig. 4) likely began to diversify 4.41 Ma in the Pliocene (2.31–6.05 Ma).

We find that for nodes 4 and 5, both molecular data and calibration ages contributed substantially to the posterior distribution of node ages (as seen in the difference between the “prior” and the “posterior” and “calibrated prior” distributions, respectively; Fig. 4); the posterior and calibrated prior distributions are strongly shifted to younger ages than the tree prior distribution. However, the posterior estimate of node 3 appears mostly influenced by the tree prior and to some extent by the calibration ages (calibrated prior). In contrast, the root age posterior differs substantially from both the tree prior and calibrated prior, suggesting that molecular data has a significant effect on the posterior distribution of Alstroemeriaceae crown age. Unsurprisingly, the Luzuriageae crown age is strongly influenced by the calibration ages (calibrated prior), as this is the node to which the fossil calibration was applied.

### Biogeographic History of Alstroemeriaceae

We infer a combined southern South American and Australasian (*PP* = 0.24), southern South American (*PP* = 0.19), or Australasian (*PP* = 0.18) origin of Alstroemeriaceae (Node 1, Fig. 5) during the Late Cretaceous. Subsequently, Alstroemerieae (Node 3, Fig. 5) moved in southern South America while Luzuriageae remained in the combined range until *Luzuriaga* and *Drymophila* diverged (Nodes 6 and 7, Fig. 5). *Bomarea* then began to move north, with the probability of an Andean ancestral range increasing through time until the ancestor of core *Bomarea* (Node 8, Fig. 5, pp = 1).

**Figure 5:**
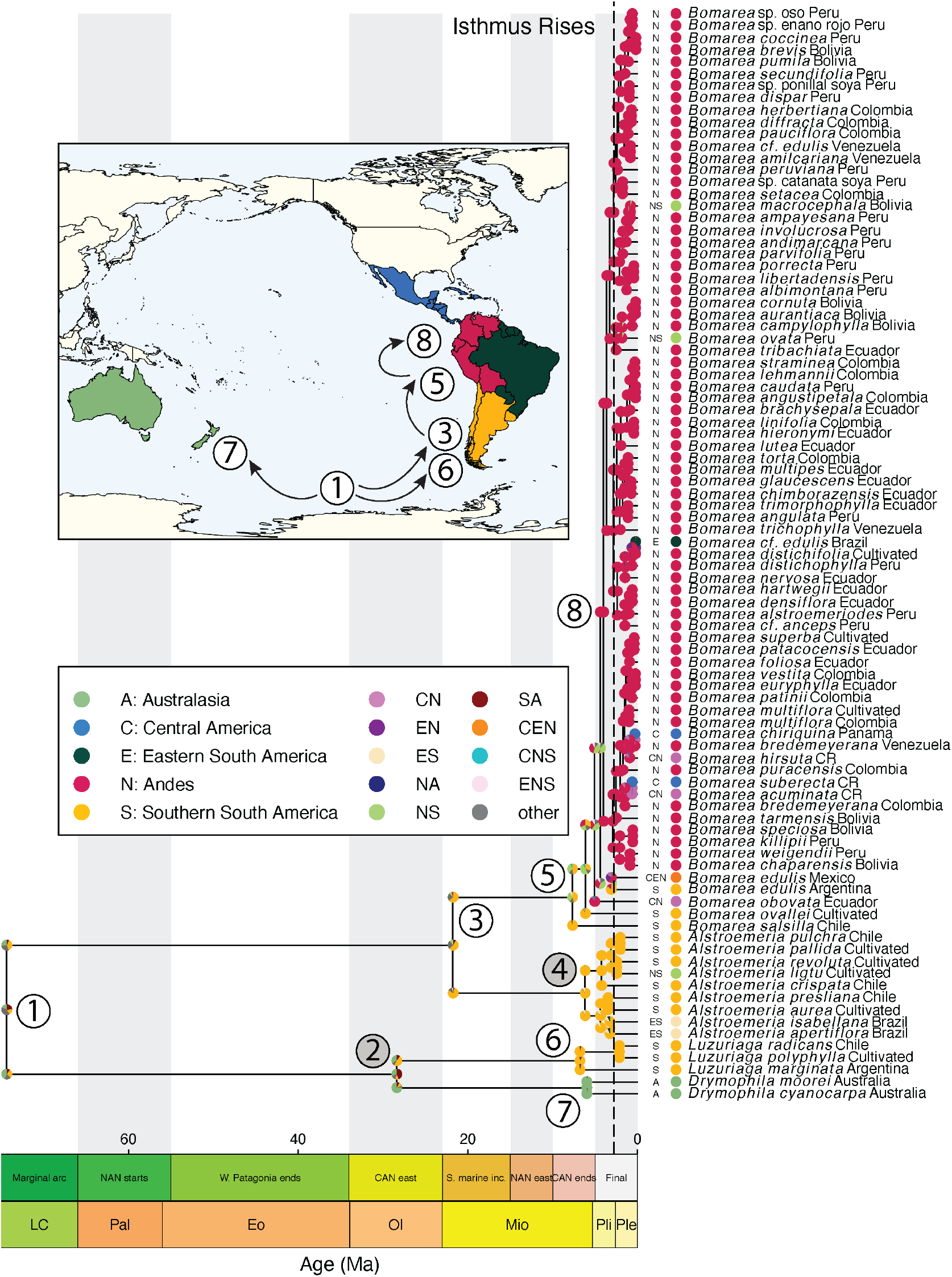
Chronogram of Alstroemeriaceae showing estimated ancestral reconstruction of biogeographic ranges. Pie charts at nodes show the top three most probably ancestral states (color of the slices) and their corresponding posterior probabilities (size of the slices). The inset map shows major biogeographic events and numbers correspond to node numbers (in white circles) labeled on the phylogeny. If node numbers are in grey circles, they are not shown on the map. The geological timescales show (top) the major events of Andean uplift (from Pérez-Escobar et al., 2022) and (bottom) the relevant geological epochs. The vertical, dashed line represents an estimate for the rise of the isthmus of Panama (2.8 myr, O’Dea et al., 2016).

### Biogeographic History of *Bomarea*

Within *Bomarea* we infer a southern South American ancestral range (*pp* = 0.38), with additional support for a combined southern South American and central Andean range (*pp* = 0.32). After *Bomarea salsilla* splits from the rest of *Bomarea*, we infer a combined range of southern South America, central Andes, northern Andes, eastern South America, and Central America, followed by a cladogenetic split in which *Bomarea ovallei* moves into southern South America and the rest of *Bomarea* moves into the central Andes. We also infer a central Andean origin of the branch subtending *Bomarea obovata* followed by range expansion into the northern Andes and Central America. Historical ranges of the *Bomarea edulis* clade are highly uncertain. We infer a combined central and southern Andean ancestor of *B. edulis* (*pp* = 0.24), with a cladogenetic split of the range into southern Andean for the Argentina with particularly high uncertainty about how *B. edulis* spread to occupy all of South and Central America.

According to our results, core *Bomarea* originated in the Andes during the Pliocene, approximately 4.41 Ma. Three clades (nodes 9, 10, and 11 in Fig. 6) then diverge around the same time in the late Pliocene. In all three clades, dispersal to and from the central and northern Andes explain most of the biogeographic reconstruction. Our results indicate that Clade 9 originated in the central Andes. A single cladogenetic dispersal event during the beginning of the Pleistocene (indicated by yellow star in Clade 9) led to the separation and subsequent diversification of a northern subclade within this lineage. Within the northern subclade, a single species (*Bomarea superba* Herb.) appears to have dispersed back to the central Andes. The phylogenetic placement of *B. superba* aligns nicely with its inferred affinities based on morphology; Hofreiter (2008) suggests that *B. superba* is more closely related to the northern Andean taxa than other Peruvian species. In addition, several lineages independently migrated further north into Central America (*e*.*g*., *Bomarea chiriquina, Bomarea suberecta*). These Central American species appear to represent a combination of independent dispersal from northern South America and in situ diversification.

**Figure 6:**
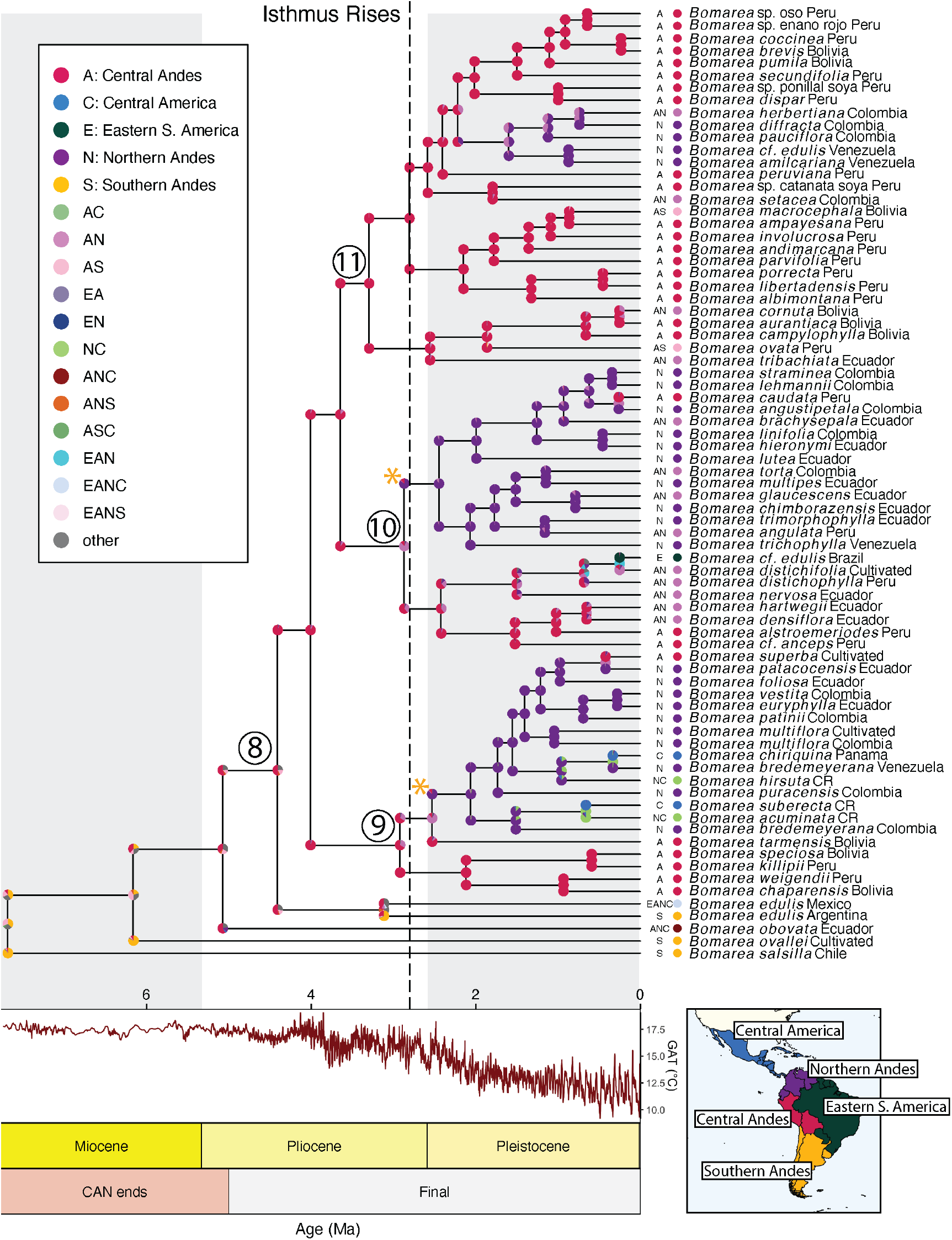
Chronogram of *Bomarea* showing estimated ancestral reconstruction of biogeographic ranges. X-axis includes (top) a reconstruction of global average temperature (GAT) over geological time (data from Tierney et al., 2020), (middle) geological epochs, and (bottom) stages of Andean uplift from Pérez-Escobar et al. (2022). Pie charts at nodes show the top three most probably ancestral states (color of the slices) and their corresponding posterior probabilities (size of the slices). Yellow stars indicate relevant cladogenetic events leading to northern Andean sub-clades. Inset map shows the five biogeographic regions.

Clade 10 originated in a combined central and northern Andean range, followed by an immediate cladogenetic range split (indicated by yellow star in Clade 10) into a primarily northern subclade and a primarily central subclade. Within the northern subclade, a single dispersal event led to the establishment of one central Andean species (*Bomarea caudata*). Many of these northern subclade species grow in high-elevation páramo habitats with an erect growth form, including *Bomarea straminea* and *Bomarea lehmanii*. These two species are morphologically similar and sister species in the phylogeny, suggesting that they may be synonyms. The central subclade served as the source of one dispersal event into Eastern South America (*Bomarea cf. edulis*) and comprises several species with contiguous northern and central Andean ranges (*e*.*g*., *Bomarea distichifolia, B. distichophylla*).

Our results show that Clade 11 originated in the central Andes, where the majority of extant species within this lineage are still distributed. Within this lineage, there is a single anagenetic dispersal event leading to a northern Andean subclade (*e*.*g*., *Bomarea diffracta*) and several species with ranges that extend over multiple defined areas, either extending to include the central and northern Andes or the central and southern Andes. *Bomarea pauciflora* is also part of Clade 11, despite its strong morphological affinities to the erect and suberect species in clade 10 (*e*.*g*., *B. angustipetala*).

### Diversification-Rate Estimation

Diversification rates in *Bomarea* increase in the core *Bomarea* clade (node 8)—the average rate in that clade is 0.78 events per lineage per million years (ELMyrs) versus 0.36 ELMyrs in the rest of the tree (Fig. 7). Rates are generally slower along the terminal branches of species in the grade (*B. salsilla, B. ovallei, B. obovata*, and *B. edulis*). We find that, on average, net-diversification rates are higher at Andean (0.74 ELMyrs) vs. non-Andean (0.665 ELMyrs) nodes, with moderate statistical support (2lnBF = 3.51).

**Figure 7:**
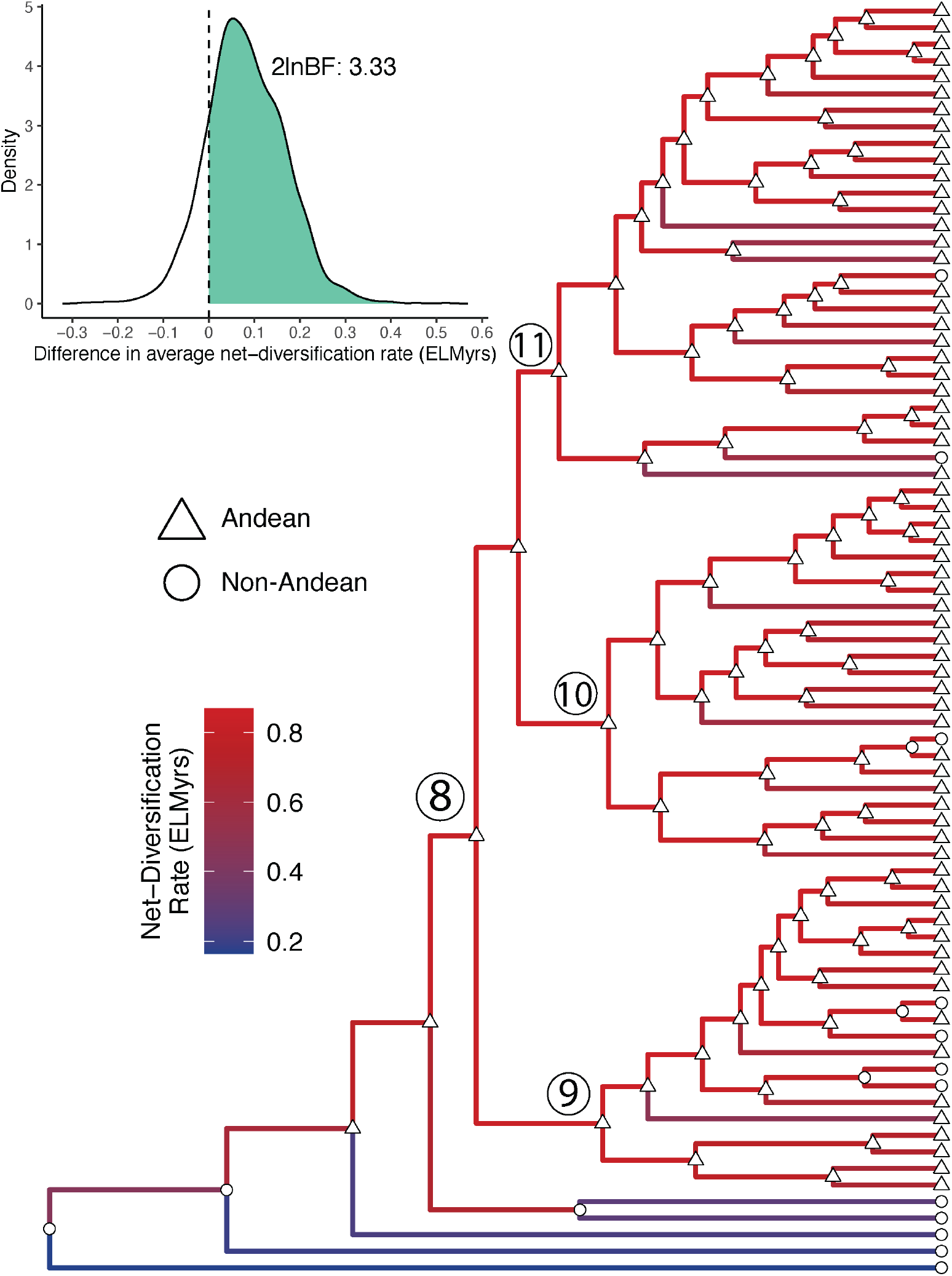
Branch-specific diversification rate estimation. The *Bomarea*-only chronogram has branches colored by estimated net-diversification rate. Node symbols represent if the node was estimated as Andean (northern or central Andean) or not in the above-described biogeographic analysis. The density plot shows the support for the average difference in net-diversification rate between Andean and non-Andean nodes in the tree, while accounting for uncertainty in the geographic state and branch rate estimates.

## Discussion

### Geological and climatic drivers of diversification and biogeography

Our inference that Andean orogeny shaped both the north-to-south biogeographic history and the rapid diversification of *Bomarea* adds to a growing body of scholarship that links geological processes to biodiversity generation in the Americas. Current evidence suggests that fewer Andean plant lineages followed the south-to-north pattern of colonization than in-situ or north-to-south (Bacon et al., 2018). However, those groups that did spread south-to-north—such as *Puya* (Jabaily and Sytsma, 2013), wax palms (Sanén et al., 2016), *Chuquiraga* (Ezcurra, 2002), *Gunnera* (Bacon et al., 2018), and *Bomarea*—tracked the rise of the Andes and adapted to new habitats as they formed. Thus, we expect—and demonstrate—that orogenic events directly shaped how and when *Bomarea* spread north and diversified.

During *Bomarea*’s origin in the mid-Miocene, Andean orogenetic activity was concentrated in the Central Andes. By the mid-Pliocene, higher elevation habitats in the central Andes were well established. The climate had also begun to cool down from the middle Miocene Climatic Optimum, perhaps facilitating the evolution of colder-tolerant groups in the tropics, including those adapted to higher elevation tropical habitats such as cloud forests and alpine regions. Core-*Bomarea* diverged from the rest of the genus—and began to diversify in the central Andes—in the context of these environmental conditions: well-established central-Andean habitats and cooling temperatures

During the final stages of Andean uplift—which were concentrated in the northern Andes—*Bomarea* began to move into and diversify in that region. Within core *Bomarea*, we observe a general nested pattern of northward dispersal, with taxa dispersing from the central Andes, diversifying, and then dispersing farther north (or, rarely, to the east). The three main clades within core *Bomarea* (indicated by nodes 9, 10, and 11 in Fig. 6) originated in the central Andes or a widespread range that includes the central Andes. These clades then served as the source of multiple independent dispersal events to the northern Andes that, through subsequent diversification, led to the establishment of extant diversity in the northern Andes. In some cases, the central Andean region also served as the source of dispersal back south, leading to the establishment of species such as *Bomarea macrocephala* and *Bomarea ovata*, whose ranges extend into northern Argentina. The clades that established in the northern Andes then served as sources of later dispersal further north in Central America and, in one case, east into Brazil.

Central American *Bomarea* are highly phylogenetically clustered. Of the six Central American species included in our phylogeny, two are restricted to central America, two are distributed across Central America and the northern Andes, and two are widely distributed species (*B. edulis* and *B. obovata*) with a range that includes Central America. Apart from the two widely distributed species, all Central American taxa are placed in the northern South American subclade of clade 9. However, they appear to represent three independent dispersal events. First, *Bomarea chiriquina* appears to have moved into Central America from a widespread ancestor through cladogenetic range splitting—the other sister lineage stayed in northern South America (represented here by a Venezuelan *B. bredemeyerana* accession). Second, *Bomarea hirsuta* extended an ancestral northern South American range into Central America anagenetically. Third, the ancestor of *B. suberecta* and *B. acuminata* extended from a northern Andean range to a combined northern Andean and Central American range, which *B. acuminata* inherited while *B. suberecta* split into only Central America. This pattern indicates a possible predisposition of species from Clade 9 to migrate to and successfully establish in Central America. Further studies would be needed to investigate dispersal potential within this clade or physiological traits that might be associated with northern migration leading to establishment in Central America.

The rise of the isthmus of Panama occurred substantially (∼2.3 million years or more) before the estimated dispersal events to Central America (O’Dea et al., 2016). We thus do not find evidence that the lack justy any terrestrial corridor directly limited *Bomarea*’s northward dispersal. Nonetheless, habitats available following the emergence of land may have continued to be inhospitable to a primarily montane lineage such as *Bomarea* for some time following the closure of a seaway passage. In fact, sea level dropped substantially from ∼3 mya to ∼1 mya, with relatively little change from 1 mya to present (Fig. 1 from O’Dea et al., 2016), around when we first infer *Bomarea* dispersal events to Central America from northern South America. It is thus likely that *Bomarea*’s repeated colonization of Central America occurred only after sufficiently high-elevation habitat formed along the isthmus.

Most splits in the phylogeny appear to occur prior to the recent and rapid fluctuations in global temperature during the late Pleistocene (Pleistocene climatic oscillations), but it is possible that these rapid shifts in temperature, and the corresponding effects on habitat connectivity in alpine regions, may have reinforced species boundaries between some recently-diverged sister taxa such as *Bomarea coccinea* and *B. brevis*, which occur sympatrically in similar Peruvian cloud forests.

*Bomarea* diversification also appears tied to Andean uplift and climatic changes in the Pliocene and Pleistocene (Fig. 6). Once *Bomarea* reached the central Andes, it began to diversify quite rapidly (Fig. 4); the 72 species in core *Bomarea* emerged over an approximately 4 million-year period. This diversification rate is comparable to the rates inferred for páramo plant lineages, which have higher diversification rates than those of groups in any other biodiversity hotspot (*Bomarea*’s net-diversification rate = 0.78; median estimate for páramo radiations = 0.73 Madriñán et al., 2013). We also show that, on average, within *Bomarea*, Andean lineages diversified faster than non-Andean ones: ancestral nodes reconstructed as Andean had higher average diversification rates (average difference = 0.08 ELMyrs), due to both the production of lineages that moved out of the Andes and to in situ diversification.

### *Bomarea edulis* monophyly

While most *Bomarea edulis* individuals form a clade, a few individuals identified as *B. edulis* fall out separately (Campbell 8900 and Bunting 4817—see below for discussion of these accessions). The monophyly of most *B. edulis* accessions indicates that the species’ strikingly wide range is not due to taxonomic misclassification. Rather, the wide range appears to be “real,” perhaps caused by extensive geographic spread with limited morphological change since *B. edulis* split with core *Bomarea*, or, more plausibly, due to human influence. While our results do not directly address the role of pre-Columbian cultivation in the evolution and spread of *B. edulis*, the monophyly of (most) of our *B. edulis* accessions provides a starting point for future work that more closely examines finer-scale processes within this species.

Two individuals identified as *B. edulis* do not form a clade with the others. The first is Campbell 8900 from western Brazil. The only *Bomarea* species previously known to occur in Brazil is *B. edulis*, which is likely why Campbell 8900 was identified as *B. edulis* despite striking morphological differences—Campbell 8900 has much larger leaves than does typical *B. edulis*. This accession may represent an undescribed taxon or the rarely collected *Bomarea ulei* Kraenzl; in the phylogeny it is nested in a clade of primarily combined central Andean and northern Andean taxa. This record thus represents a second *Bomarea* lineage to disperse to and establish in Brazil. The second accession (mis)identified as *B. edulis* is Bunting 4817 from Venezuela. This herbarium specimen has fruits but no flowers, making exact identification difficult. However, the locality and fruit morphology point to it being *Bomarea amilcariana* Stergios & Dorr, and indeed it falls out sister to *B. amilcariana* in the A-MSC phylogeny.

The *B. edulis* clade also includes a few specimens not identified as *B. edulis*. Of these, one is a specimen identified as *Bomarea dolichocarpa* Killip (Barbour 5069), and it could be a misidentified *B. edulis* as the two species are commonly confused when not in fruit (Hofreiter, 2006). Alternatively, its placement could indicate that *B. dolichocarpa* and *B. edulis* are conspecific, though the differences in their fruit shapes suggests that this is unlikely. The primarily *B. edulis* clade also includes an individual identified as a putative hybrid of *B. edulis* and *Bomarea acutifolia* (Tribble 79). None of the genetic evidence suggests a hybrid origin for this accession, so it is likely an atypical *B. edulis* rather than a hybrid.

We note that the unresolved taxonomy of many *Bomarea* species—including those brought to light in this study (*e*.*g*., *B. ulei*)—is a challenge for this type of work, as issues with synonymy and species boundaries directly affect the definition of taxonomic units for birth-death models and species ranges for biogeographic inference. We thus advocate for continued taxonomic work on this group given its potential as a model of the effect of geological events on biodiversity in the tropical Americas.

### Applying genome-scale data to recent rapid radiations

We demonstrate the effectiveness of genome-scale data for phylogenetic inference of a recent and rapid radiation, but we also emphasize the importance of careful data curation and of using analytical methods appropriate for young and rapidly evolving clades.

Target capture methods are successful at generating genome-scale data for samples preserved in silica gel for DNA extraction as well as older museum-preserved samples from herbaria (Dodsworth et al., 2019). Our Alstroemeriaceae dataset includes 74 tips from herbarium-sampled material. Herbarium samples enabled the dense taxonomic sampling in our study and allowed for the inclusion of rarely collected taxa, which provided the detail and power to resolve finer-scale biogeographic processes and diversification-rate estimates.

We found high levels of contaminant sequences and a large proportion of duplicated loci in our target capture dataset (see Results: Data Processing). Contaminant sequences can be common in such datasets, especially when derived from museum samples (Andermann et al., 2020), though we note that the levels of contamination we found were unusually high. Universal probe sets for target capture such as the GoFlag angiosperm 408 probes used in this study (Endara and Burleigh, 2022), Angiosperm 353 (Johnson et al., 2019), and ultraconserved elements (UCEs; McCormack et al., 2012) are designed to target sequences from an evolutionarily broad group of organisms and thus may be also more likely to recover contaminant sequences than clade-specific probe sets. Additionally, while the GoFlag angiosperm 408 probes (and most other universal probe sets) target single-copy nuclear genes, these regions were identified as single-copy at a broad evolutionary scale (all angiosperms), which cannot ensure that recent gene duplications have not occurred within our target group (Frost and Lagomarsino, 2021). While one option may be to design more clade-specific probe sets, this is not feasible for many researchers—including for us in this study—and may negate the potential data-sharing benefits of global probe sets (depending on the way clade-specific probe sets are generated). To address these concerns, we constructed and manually inspected gene trees for all loci used in this analysis. Our manual gene-tree inspection allowed us to identify potential contaminants, confirm sequence identity through blast (Altschul et al., 1990), and remove contaminant sequences, which were present in a significant proportion of loci (161 of 221 loci). We also used manual gene-tree inspection to carefully tease apart the causes of putatively duplicated sequences—multiple sequences from one accession recovered for one locus. These duplicated sequences may contain information about gene duplication events that pertain to the entire locus and thus simply deleting duplicated accessions is not sufficient for guaranteeing orthology. Even accessions with only one recovered copy may have experienced the duplication, with only one copy recovered due to lower sequencing coverage. This concept is similar to the “hidden paralogs” discussed in Frost and Lagomarsino (2021). The authors point out that herbarium-derived data may be more prone to differential success in recovering paralogous loci, and the lack of duplicated copies may be hiding the fact that two sequences of two species are actually paralogous rather than homologous. Thus, we chose not to simply delete multiple copies when they occurred and instead to parse apart these loci using gene trees. By manually inspecting the gene trees of loci with duplicated copies, we were able to preserve most of these loci in the analysis. In some cases, we were able to split loci into two sets of homologous regions, increasing the amount of usable data. Together, these steps removed spurious sequences from the dataset and ensured that our loci represent homologous rather than paralogous sequences.

Our results also demonstrate the importance species-tree inference over concatenation approaches when rampant ILS is suspected. For each phylogeny with more than three tips, there are certain sets of parameter values that define an “anomaly zone” where the most likely gene tree is not equivalent to the true species tree (Rannala et al., 2020). Under these parameter values, using a species tree inference method that does not account for ILS will generally infer an incorrect topology. While the prevalence of this anomaly zone is unclear (Rannala et al., 2020), short branch lengths (common in our phylogeny) make gene tree-species tree conflict more likely (Maddison, 1997). The major backbone topology of *Bomarea* differs between our concatenated maximum likelihood analysis and our A-MSC analysis in `ASTRAL-II` (Fig. 3), particularly in the relative positions of *B. obovata* and *B. edulis*. It is possible that using the ML topology would have significantly impacted our biogeographic inference, given the deep position of the changing node in the tree.

### Divergence time estimation uncertainty in macroevolutionary inferences

Our approach emphasizes the importance of careful examination of node-based calibrations for divergence time estimation. The influence of prior specification on divergence time estimation has been well established (*e*.*g*.,, May et al., 2021), and previous studies have well-demonstrated issues with insufficient data for updating node-age priors (Brown and Smith, 2018). Thus, we examined the relative contributions of the tree prior and the temporal (fossil and secondary calibrations) and molecular data to the posterior node age estimates. Molecular data and secondary and fossil calibrations all contribute to our node age estimates, but the degree to which these sources of information update the tree prior varies depending on the node in question (Fig. 4). For example, our root age estimate is highly influenced by molecular data while the crown age of Alstroemerieae appears to be largely influenced by the tree prior. The ages of many nodes of the tree, especially the root and Luzuriageae and Alstroemerieae MRCAs, are highly uncertain. This analysis demonstrated the relatively weak influence of our data on the divergence times of some nodes, which gave us further impetus to incorporate temporal uncertainty in our biogeographic inferences.

## Conclusions

In this study we produce the first well-sampled phylogeny of *Bomarea*, a charismatic tropical American plant clade, and demonstrate how major geographic and climatological events of the past 80 million years have shaped its colonization of new habitats and its diversification. In particular, *Bomarea*’s origin in southern South America and its subsequent movement north as the Andes arose demonstrates how southern-temperate lineages have contributed to the outstanding diversity of tropical biomes today. We also illustrate how genome-scale data, museum-sampled material, and careful bioinformatic processing can expand our ability to infer evolutionary relationships and macroevolutionary processes. For understudied (often tropical) lineages, the steps we demonstrate allow for rigorous analyses in the face of fewer genomic resources and a dearth of well-preserved material, and thus this study may serve as a model for further work on tropical biodiversity generation.

## Author Contributions

CMT designed and executed the study. CDS and CJR assistant with project design. CMT, FA-G, EG, and AV collected data. CMT and JGB performed analyses. CMT, FA-G, JGB, RZ-F, CDS, and CJR guided interpretation of results.

## Acknowledgements

Fieldwork and sequencing costs were funded by grants from the American Society of Plant Taxonomists, the Pacific Bulb Society, the Garden Club of America, the Society of Systematic Biologists, The American Philosophical Society, the Torrey Botanical Society, the Tinker Foundation, and the Integrative Biology Department at UC Berkeley. CMT was supported by an NSF GRFP and NSF PRFB during this project.

The following herbaria graciously provided access to collections for destructive sampling: The Missouri Botanical Garden (MO), The New York Botanical Garden (NY), The Field Museum (F), the United States National Herbarium (SI), the University Herbarium (UC), the Herbario Nacional de México (MEX), and the National Herbarium of Victoria (MEL). The UC Botanical Garden, the San Francisco Botanical garden, and the Royal Botanic Gardens, Kew kindly provided access to living collections.

Marko Gomez, Victoria Sosa, Anahí Espinoza Jiménez, JosuéLuna, Rafaél Torres, Luis Gerardo Hernandez Sandova, Yolanda Pantoja, Beatriz Velásquez, Don Ermelando, Carlos Durán, Pedro de la Cruz Salvador, JoséMaría de Jesús-Almirante, Oscar Dorado, Karime Díaz, Gerardo Cuevas, Jair Esteban López Reyes, Francisco Javier Ortiz Gorostieta, Daniela Gutijr, Enrique Florentino Lopez, Herbert Jassin Sarrazola Yepes, Carlos Felipe Bolaños-Sibaja, Pablo Gallego, and Jairo Hidalgo asssisted with collections in the field.

We thank Michael R. May, Jenna B. Ekwealor, and the rest of the Rothfels lab sensu lato for their useful feedback on the manuscript.

