## Supplemental Materials for "The rapid radiation of *Bomarea* (Alstroemeriaceae: Liliales), driven by the rise of the Andes"

### Supplemental Material

#### Supplemental Section 1 Methods

##### Supplemental Section 1.1 Sequencing methods

Extractions were treated with a bead DNA cleanup and quantified. Any samples with $> 5ng$ total DNA were discarded. All remaining samples were sheared to 300 base pairs (bp) average size prior to library prep. Library preparation was then performed with these sheared extractions by repairing fragment ends and adding adenine residue to blunt-end fragments on the 3′ end (Bentley et al., 2008). Libraries were then ligated to Illumina-appropriate barcoded adapters and amplified with PCR for 9-11 cycles.

#### Supplemental Section 2 Specimen data

**Table S1:** Voucher information for specimens used in phylogenetic reconstruction.

| | Collection Information | | | | Included | | |
|---|---|---|---|---|---|---|---|
| **Scientific Name** | **Number** | **Region** | **Material** | **Institution** | **Sequenced** | **In phylogram** | **In chronogram** |
| *Alstroemeria apertiflora* | Hatschbach 17552 | Brazil | herb. (1967) | UC | Y | Y | Y |
| *Alstroemeria aurea* | Andrews 94004 | Cultivated | silica | UCBG | Y | Y | Y |
| *Alstroemeria crispata* | Wegenknecht 18128 | Chile | herb. (1940) | UC | Y | Y | Y |
| *Alstroemeria haemantha* | UCBG54.397 | Chile | silica | UC | Y | | |
| *Alstroemeria inodora* | Irwin 2703 | Brazil | herb. (1959) | UC | Y | | |
| *Alstroemeria isabellana* | Hatschbach 43366 | Brazil | herb. (1980) | UC | Y | Y | Y |
| *Alstroemeria ligtu* | Ornduff 9112 | Cultivated | silica | UCBG | Y | Y | Y |
| *Alstroemeria nervosa* | Hatschbach 34050 | Brazil | herb. (1974) | UC | | | |
| *Alstroemeria pallida* | Chase 19989 | Cultivated | living | K | Y | Y | Y |
| *Alstroemeria presliana* | Henchie 131 | Chile | silica | K | Y | Y | Y |
| *Alstroemeria pulchra* | Kelch 00.014 | Chile | herb. (2000) | UC | Y | Y | Y |
| *Alstroemeria pygmaea* | Saunders 715 | Unknown | unknown | K | | | |
| *Alstroemeria revoluta* | Watson 6608 | Cultivated | herb. (2019) | UCBG | Y | Y | Y |
| *Alstroemeria stenosepala* | Hatschbach 38142 | Brazil | herb. (1976) | UC | | | |
| *Bomarea acuminata* | Bonifacino 6051 | CR | silica | CR | Y | Y | Y |
| *Bomarea acutifolia* | Tribble 79 | Mexico | silica | UC | Y | Y | |
| *Bomarea aff. cruenta* | Diaz 3785 | Peru | herb. (1989) | MO | Y | | |
| *Bomarea albimontana* | Leiva S.N. | Peru | herb. (2002) | F | Y | Y | Y |
| *Bomarea allenii* | D'Arcy 12440 | Panama | herb. (1979) | MO | Y | | |
| *Bomarea alstroemeriodes* | Dillon 1747 | Peru | herb. (1979) | F | Y | Y | Y |
| *Bomarea amazonica* | Barbour 2859 | Peru | herb. (1978) | MO | | | |
| *Bomarea amilcariana* | Dorr 8337 | Venezuela | herb. (1998) | F | Y | Y | Y |
| *Bomarea ampayesana* | Bocke 3215 | Peru | herb. (1980) | NY | Y | Y | Y |
| *Bomarea andimarcana* | Suclli 1307 | Peru | herb. (2003) | MO | Y | Y | Y |
| *Bomarea andreana* | Stein 3218A | Colombia | herb. (1986) | MO | Y | | |
| *Bomarea angulata* | Quiroz 3208 | Peru | herb. (1992) | MO | Y | Y | Y |
| *Bomarea angustipetala* | Alzate 5116 | Colombia | silica | HUA | Y | Y | Y |
| *Bomarea angustissima* | Macbride 4409 | Peru | herb. (1923) | F | Y | | |
| *Bomarea aurantiaca* | Fuentes 13880 | Bolivia | herb. (2009) | MO | Y | Y | Y |
| *Bomarea boliviensis* | Eyerdam 24757 | Bolivia | herb. (1939) | F | Y | | |
| *Bomarea brachysepala* | Jorgensen 2129 | Ecuador | herb. (2000) | MO | Y | Y | Y |
| *Bomarea bracteolata* | McPherson 10126 | Panama | herb. (1986) | MO | Y | | |
| *Bomarea bredemeyerana* | Tribble 06 | Colombia | silica | UC | Y | Y | Y |
| *Bomarea bredemeyerana* | Liesner 7935 | Venezuela | herb. (1979) | MO | Y | Y | Y |
| *Bomarea brevis* | Teran 1759 | Bolivia | herb. (2007) | MO | Y | Y | Y |
| *Bomarea campanularia* | Vargas 5527 | Ecuador | herb. (2005) | MO | Y | | |
| *Bomarea campylophylla* | Maldonado 3174 | Bolivia | herb. (2002) | MO | Y | Y | Y |
| *Bomarea carderi* | Devia 1160 | Colombia | herb. (1986) | MO | | | |
| *Bomarea caucana* | Arbelaez 6201 | Colombia | herb. (1939) | US | Y | | |
| *Bomarea caudata* | Weberbauer 7559 | Peru | herb. (1926) | F | Y | Y | Y |
| *Bomarea caudatisepala* | Hammel 7396 | Panama | herb. (1979) | MO | | | |
| *Bomarea ceratophora* | Dodson 14168 | Ecuador | herb. (1984) | MO | | | |
| *Bomarea cf. anceps* | Lopez M. 8913 | Peru | herb. (1981) | MO | Y | Y | Y |
| *Bomarea chaparensis* | Steinbach 8897 | Bolivia | herb. (1929) | MO | Y | Y | Y |
| *Bomarea chimborazensis* | Aedo 13023 | Ecuador | herb. (2006) | MO | Y | Y | Y |
| *Bomarea chiriquina* | Hammel 6374 | Panama | herb. (1979) | MO | Y | Y | Y |
| *Bomarea coccinea* | Nunez 13229 | Peru | herb. (1991) | MO | Y | Y | Y |
| *Bomarea colombiana* | Cuatrecasas 1959 | Colombia | herb. (1959) | US | Y | | |
| *Bomarea cordifolia* | Foster 7709 | Peru | herb. (1984) | MO | Y | | |
| *Bomarea cornigera* | Sanchez Vega 3692 | Peru | herb. (1985) | MO | Y | | |
| *Bomarea cornuta* | Fuentes 17664 | Bolivia | herb. (2012) | MO | Y | Y | Y |
| *Bomarea costaricensis* | Hammel 20590 | CR | herb. (1996) | MO | Y | | |
| *Bomarea crassifolia* | Betancur 2988 | Colombia | herb. (1993) | MO | Y | | |
| *Bomarea crocea* | Galiano 6868 | Peru | herb. (2004) | MO | | | |
| *Bomarea densiflora* | Alzate 3151 | Ecuador | silica | QCNE | Y | Y | Y |

**Table S1:** Voucher information for specimens used in phylogenetic reconstruction. *(continued)*

| | Collection Information | | | | Included | | |
|---|---|---|---|---|---|---|---|
| **Scientific Name** | **Number** | **Region** | **Material** | **Institution** | **Sequenced** | **In phylogram** | **In chronogram** |
| *Bomarea denticulata* | Plowman S.N. | Peru | herb. (1981) | F | Y | | |
| *Bomarea diffracta* | Tribble 12 | Colombia | silica | HUA | Y | Y | Y |
| *Bomarea dispar* | Schunke V. 3544 | Peru | herb. (1969) | F | Y | Y | Y |
| *Bomarea dissitifolia* | Paniagua 8786 | Peru | herb. (2012) | MO | Y | | |
| *Bomarea distichifolia* | SFBG2012-0053 | Cultivated | silica | SFBG | Y | Y | Y |
| *Bomarea distichophylla* | Rojas 2689 | Peru | herb. (2004) | F | Y | Y | Y |
| *Bomarea dolichocarpa* | Barbour 5069 | Peru | herb. (1980) | F | Y | Y | |
| *Bomarea dolichocarpa* | Phillips 110 | Peru | herb. (1989) | MO | Y | | |
| *Bomarea dulcis* | Fuentes 12519 | Bolivia | herb. (2008) | MO | | | |
| *Bomarea edulis* | Deginani 1587 | Argentina | herb. (2000) | MO | Y | Y | |
| *Bomarea edulis* | Hunziker 12327 | Argentina | herb. (1992) | MO | Y | Y | |
| *Bomarea edulis* | Múlgura de Romero 1876 | Argentina | herb. (1997) | MO | Y | Y | |
| *Bomarea edulis* | Novara 2378 | Argentina | herb. (1982) | MO | Y | Y | Y |
| *Bomarea edulis* | Porta 193 | Argentina | herb. (1943) | MO | | | |
| *Bomarea edulis* | Zuloaga 5899 | Argentina | herb. (1997) | MO | Y | | |
| *Bomarea edulis* | Mamani 458 | Bolivia | herb. (1995) | MO | Y | | |
| *Bomarea edulis* | Solomon 7616 | Bolivia | herb. (1982) | MO | Y | Y | |
| *Bomarea edulis* | Campbell 8900 | Brazil | herb. (1986) | MO | Y | Y | Y |
| *Bomarea edulis* | Thomas 13631 | Brazil | herb. (2004) | NY | Y | Y | |
| *Bomarea edulis* | Irwin 12289 | Brazil | herb. (1966) | NY | Y | | |
| *Bomarea edulis* | Pires 6119 | Brazil | herb. (1956) | NY | Y | | |
| *Bomarea edulis* | Melo 06 | Brazil | herb. (2004) | NY | Y | | |
| *Bomarea edulis* | Eiten 10407 | Brazil | herb. (1970) | US | Y | | |
| *Bomarea edulis* | Kegler 89 | Brazil | herb. (1999) | US | Y | Y | |
| *Bomarea edulis* | Eiten 9877 | Brazil | herb. (1968) | US | Y | Y | |
| *Bomarea edulis* | Duśen 14398 | Brazil | herb. (1914) | US | Y | | |
| *Bomarea edulis* | Fontella 101 | Brazil | herb. (1961) | US | Y | | |
| *Bomarea edulis* | Gentry 47615 | Colombia | herb. (1984) | MO | Y | | |
| *Bomarea edulis* | Madriñón 506 | Colombia | herb. (1989) | MO | Y | | |
| *Bomarea edulis* | J.F. Morales 6635 | CR | herb. (1998) | MO | Y | Y | |
| *Bomarea edulis* | Sanchez S.N. | Cuba | herb. (1991) | HAJB | Y | | |
| *Bomarea edulis* | Smith 3264 | Cuba | herb. (1936) | US | Y | Y | |
| *Bomarea edulis* | Jimenez 3903 | DR | herb. (1958) | US | Y | Y | |
| *Bomarea edulis* | Breckon 2075 | Guatemala | herb. (1976) | F | Y | Y | |
| *Bomarea edulis* | Jansen-Jacobs 4459 | Guyana | herb. (1995) | MO | Y | | |
| *Bomarea edulis* | Mass, P.S.M. 7186 | Guyana | herb. (1988) | NY | | | |
| *Bomarea edulis* | Leonard 9424 | Haiti | herb. (1929) | US | Y | | |
| *Bomarea edulis* | Proctor 10821 | Haiti | herb. (1955) | US | Y | Y | |
| *Bomarea edulis* | Harmon 3784 | Honduras | herb. (1970) | MO | Y | Y | |
| *Bomarea edulis* | Tribble 90 | Mexico | silica | MEXU | Y | Y | |
| *Bomarea edulis* | Tribble 32-5 | Mexico | silica | MEXU | Y | Y | Y |
| *Bomarea edulis* | Tribble 37-7 | Mexico | silica | MEXU | Y | Y | |
| *Bomarea edulis* | Tribble 113 | Mexico | silica | MEXU | Y | Y | |
| *Bomarea edulis* | Tribble 76 | Mexico | silica | MEXU | Y | Y | |
| *Bomarea edulis* | Tribble 70-1 | Mexico | silica | MEXU | Y | Y | |
| *Bomarea edulis* | Stevens 21839 | Nicaragua | herb. (1982) | MO | Y | Y | |
| *Bomarea edulis* | Duke 13712 | Panama | herb. (1967) | MO | Y | Y | |
| *Bomarea edulis* | Evrard 9623 | Peru | herb. (1982) | MO | | | |
| *Bomarea edulis* | Vasquez 20845 | Peru | herb. (1996) | MO | Y | | |
| *Bomarea edulis* | Herrera 10093 | Suriname | herb. (2003) | MO | | Y | |
| *Bomarea edulis* | Lindeman 455 | Suriname | herb. (1988) | NY | Y | | |
| *Bomarea edulis* | Aymard C. 1314 | Venezuela | herb. (1892) | MO | Y | Y | |
| *Bomarea edulis* | Gonzalez 228 | Venezuela | herb. (1979) | F | Y | | |
| *Bomarea edulis* | Bunting 4817 | Venezuela | herb. (1975) | NY | Y | Y | Y |
| *Bomarea edulis* | Tribble 55-2 | Veracruz | silica | MEXU | Y | Y | |
| *Bomarea edulis* | Tribble 65 | Veracruz | silica | MEXU | Y | Y | |
| *Bomarea euryphylla* | Vargas 2930 | Ecuador | herb. (1998) | MO | Y | Y | Y |
| *Bomarea fimbriata* | Ledezma N. 710 | Bolivia | herb. (2004) | MO | Y | | |
| *Bomarea foliosa* | Zak 2268 | Ecuador | herb. (1987) | NY | Y | Y | Y |
| *Bomarea formosissima* | Vasquez 32895 | Peru | herb. (2007) | MO | Y | | |
| *Bomarea glaucescens* | Caranqui 1583 | Ecuador | herb. (2006) | MO | Y | Y | Y |
| *Bomarea goniocaulon* | Paniagua Z. 8753 | Peru | herb. (2012) | MO | | | |
| *Bomarea graminifolia* | Vargas 306 | Ecuador | herb. (1995) | MO | Y | | |
| *Bomarea hartwegii* | Alzate 3157 | Ecuador | silica | QCNE | Y | Y | Y |
| *Bomarea herbertiana* | Pennell 3224 | Colombia | herb. (1917) | NY | Y | Y | Y |
| *Bomarea hieronymi* | Alvarez 2557 | Ecuador | herb. (2000) | MO | Y | Y | Y |
| *Bomarea hirsuta* | Croat 69997 | Colombia | herb. (1990) | MO | | | |
| *Bomarea hirsuta* | Bonifacino 6052 | CR | silica | CR | Y | Y | Y |
| *Bomarea huanuco* | Diaz S. 2249 | Peru | herb. (1987) | MO | Y | | |
| *Bomarea involucrosa* | Galiano 6988 | Peru | herb. (2004) | MO | Y | Y | Y |
| *Bomarea killipii* | Vasquez 33143 | Peru | herb. (2007) | MO | Y | Y | Y |
| *Bomarea lancifolia* | Jorgensen 1074 | Ecuador | herb. (1994) | MO | Y | | |
| *Bomarea lehmannii* | Alzate 3205 | Colombia | silica | HUA | Y | Y | Y |
| *Bomarea libertadensis* | Alayo 020 | Peru | herb. (1988) | F | Y | Y | Y |
| *Bomarea linifolia* | Tribble 8 | Colombia | silica | HUA | Y | Y | Y |
| *Bomarea longipes* | Neill 15360 | Ecuador | herb. (2006) | MO | Y | | |
| *Bomarea longistyla* | Vilcapoma 1730 | Peru | herb. (1992) | MO | Y | | |
| *Bomarea lopezii* | Rodriguez 3125 | Peru | herb. (2006) | MO | Y | | |
| *Bomarea lutea* | van der Werff 12169 | Ecuador | herb. (1991) | MO | Y | Y | Y |
| *Bomarea macrocephala* | Vargas 7052 | Bolivia | herb. (2003) | MO | Y | Y | Y |
| *Bomarea macusani* | Nunez V. 8477 | Peru | herb. (1987) | MO | Y | | |
| *Bomarea moritziana* | Palacios 11972 | Ecuador | herb. (2006) | MO | | | |
| *Bomarea multiflora* | Alzate 4974 | Colombia | silica | HUA | Y | Y | Y |
| *Bomarea multiflora* | Greenhouse | Cultivated | silica | UC | Y | Y | Y |
| *Bomarea multipes* | Alzate 3153 | Ecuador | silica | QCNE | Y | Y | Y |
| *Bomarea nematocaulon* | Foster 10534 | Peru | herb. (1985) | F | Y | | |
| *Bomarea nervosa* | Alzate 3154 | Ecuador | silica | QCNE | Y | Y | Y |
| *Bomarea obovata* | Bonifacino 6050 | CR | silica | CR | Y | Y | |
| *Bomarea obovata* | Clark 4985 | Ecuador | herb. (1998) | MO | Y | Y | Y |
| *Bomarea ovallei* | Chase 521 | Cultivated | living | K | Y | Y | Y |
| *Bomarea ovata* | Farfan 526 | Peru | herb. (2004) | MO | Y | Y | Y |
| *Bomarea ovata* | Ugent 3791 | Peru | herb. (1963) | UC | Y | | |
| *Bomarea ovata* | Agostini 2763 | Venezuela | herb. (1983) | MO | Y | | |
| *Bomarea pardina* | Croat 94359 | Ecuador | herb. (2005) | MO | Y | | |
| *Bomarea parvifolia* | Stein 2019 | Peru | herb. (1985) | MO | Y | Y | Y |
| *Bomarea patacocensis* | Luteyn 14078 | Ecuador | herb. (1990) | MO | Y | Y | Y |
| *Bomarea patinii* | Tribble 3 | Colombia | silica | HUA | Y | Y | Y |
| *Bomarea pauciflora* | Alzate 2925 | Colombia | silica | HUA | Y | Y | Y |
| *Bomarea perglabra* | Lojtnant A. 13708 | Ecuador | herb. (1979) | MO | Y | | |
| *Bomarea peruviana* | Stein 2033B | Peru | herb. (1985) | MO | Y | Y | Y |
| *Bomarea phyllostachya* | Leiva 1768 | Peru | herb. (1996) | MO | | | |
| *Bomarea porrecta* | Alayo B. 020 | Peru | herb. (1988) | MO | Y | Y | Y |
| *Bomarea pumila* | Fuentes 13830 | Bolivia | herb. (2009) | MO | Y | Y | Y |

**Table S1:** Voucher information for specimens used in phylogenetic reconstruction. *(continued)*

| | Collection Information | | | | Included | | |
|---|---|---|---|---|---|---|---|
| **Scientific Name** | **Number** | **Region** | **Material** | **Institution** | **Sequenced** | **In phylogram** | **In chronogram** |
| *Bomarea puracensis* | Garcia-Barriga 13025 | Colombia | herb. (1948) | US | Y | Y | Y |
| *Bomarea purpurea* | Leiva 929 | Peru | herb. (1993) | MO | Y | | |
| *Bomarea rosea* | Smith 3541 | Peru | herb. (1983) | MO | | | |
| *Bomarea salsilla* | Ackerman 545 | Chile | herb. (2002) | MO | Y | Y | Y |
| *Bomarea sclerophylla* | Smith 7731 | Peru | herb. (1984) | MO | Y | | |
| *Bomarea secundifolia* | Macbride 4962 | Peru | herb. (1923) | F | Y | Y | Y |
| *Bomarea setacea* | Tribble 7 | Colombia | silica | HUA | Y | Y | Y |
| *Bomarea sp. 'catanata soya'* | Graham 12611 | Peru | silica | F | Y | Y | Y |
| *Bomarea sp. 'enano rojo'* | Graham 12599 | Peru | silica | F | Y | Y | |
| *Bomarea sp. 'enano rojo'* | Graham 12607 | Peru | silica | F | Y | Y | Y |
| *Bomarea sp. 'enano verde'* | Graham 12600 | Peru | silica | F | Y | Y | |
| *Bomarea sp. 'enano verde'* | Graham 12609 | Peru | silica | F | Y | Y | |
| *Bomarea sp. 'oso'* | Graham 12613 | Peru | silica | F | Y | Y | Y |
| *Bomarea sp. 'ponillal soya'* | Graham 12616 | Peru | silica | F | Y | Y | Y |
| *Bomarea speciosa* | Cornejo 800 | Bolivia | herb. (2009) | MO | Y | Y | Y |
| *Bomarea straminea* | Alzate 3300 | Colombia | silica | HUA | Y | Y | Y |
| *Bomarea suberecta* | Bonifacino 6053 | CR | silica | CR | Y | Y | Y |
| *Bomarea superba* | SFBG2012-0052*A | Cultivated | silica | SFBG | Y | Y | Y |
| *Bomarea tarmensis* | Fernandez 3467 | Bolivia | herb. (2005) | MO | Y | Y | Y |
| *Bomarea torta* | Alzate 3270 | Colombia | silica | HUA | Y | Y | Y |
| *Bomarea tribachiata* | Jorgensen 158 | Ecuador | herb. (1994) | MO | Y | Y | Y |
| *Bomarea trichophylla* | Rivero 1217 | Venezuela | herb. (1987) | MO | Y | Y | Y |
| *Bomarea trimorphophylla* | Alzate 3158 | Ecuador | silica | QCNE | Y | Y | Y |
| *Bomarea uncifolia* | Jorgensen 931 | Ecuador | herb. (1994) | MO | Y | | |
| *Bomarea uniflora* | Huamantupa 8349 | Peru | herb. (2006) | MO | Y | | |
| *Bomarea velscoana* | Vargas 11109 | Peru | herb. (1939) | F | Y | | |
| *Bomarea vestita* | Alzate 2814 | Colombia | silica | HUA | Y | Y | Y |
| *Bomarea vitellina* | Stein 3467 | Colombia | herb. (1986) | MO | Y | | |
| *Bomarea weigendii* | Davis 1224 | Peru | herb. (1981) | F | Y | Y | Y |
| *Drymophila cyanocarpa* | Messina 989 | Australia | herb. (2016) | MEL | Y | Y | Y |
| *Drymophila moorei* | Copeland 4560 | Australia | silica | NSW | Y | Y | Y |
| *Luzuriaga marginata* | Bonifacino 550 | Argentina | herb. (2002) | US | Y | Y | Y |
| *Luzuriaga polyphylla* | UCBG90.2401 | Cultivated | silica | UCBG | Y | Y | Y |
| *Luzuriaga radicans* | Layton 84 | Chile | herb. (2008) | US | Y | Y | Y |

#### Supplemental Section 3 Phylogenetic inference

##### Supplemental Section 3.1 ASTRAL-II inference

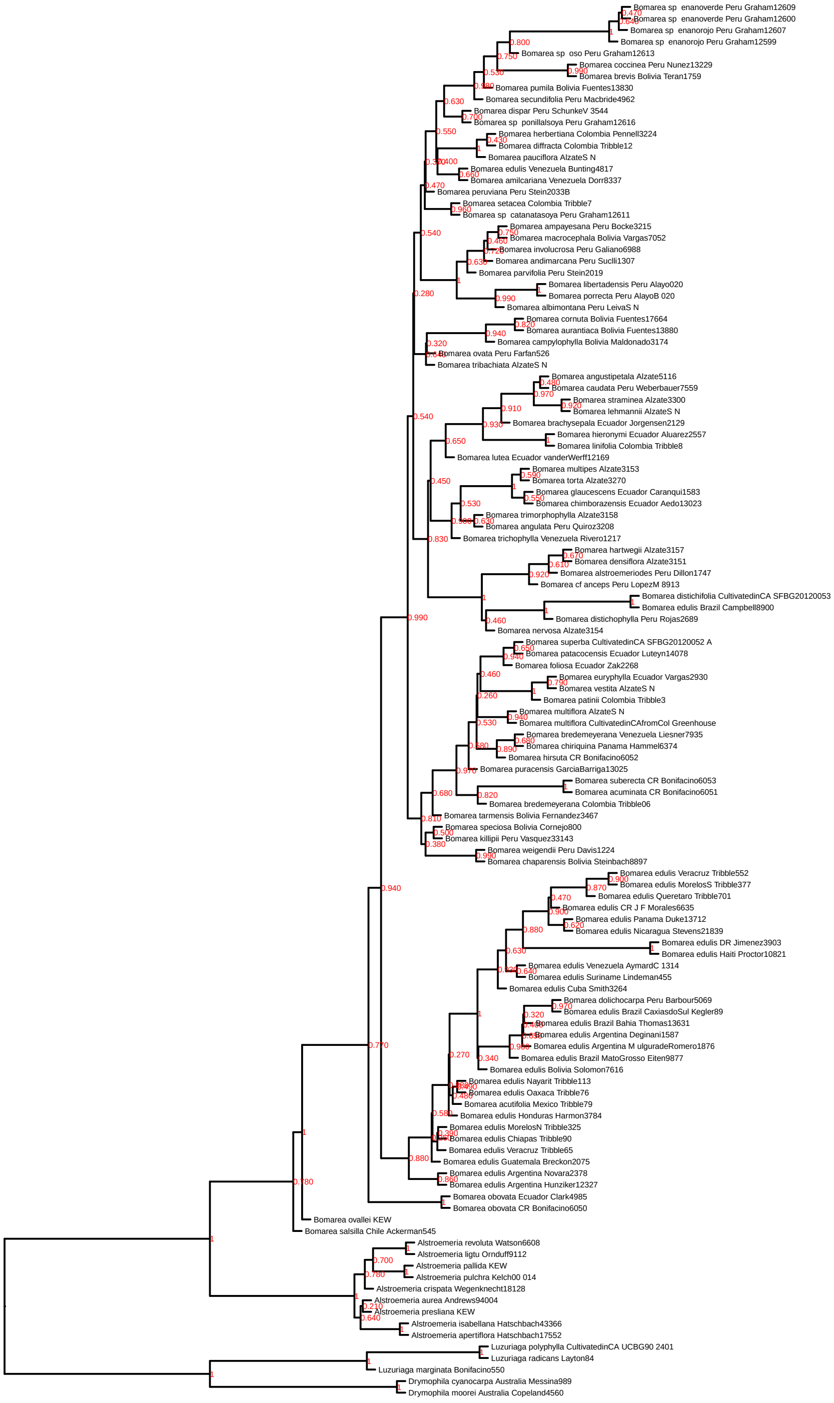


**Figure S1:** Phylogeny produced by ASTRAL-III analysis. Because each tip in the analysis is only one individual, ASTRAL cannot infer the lengths of terminal branches. We artificially set those branch lengths to be 0.1 for visualization purposes.

#### Supplemental Section 3.2 Maximum likelihood inference with `IQtree`

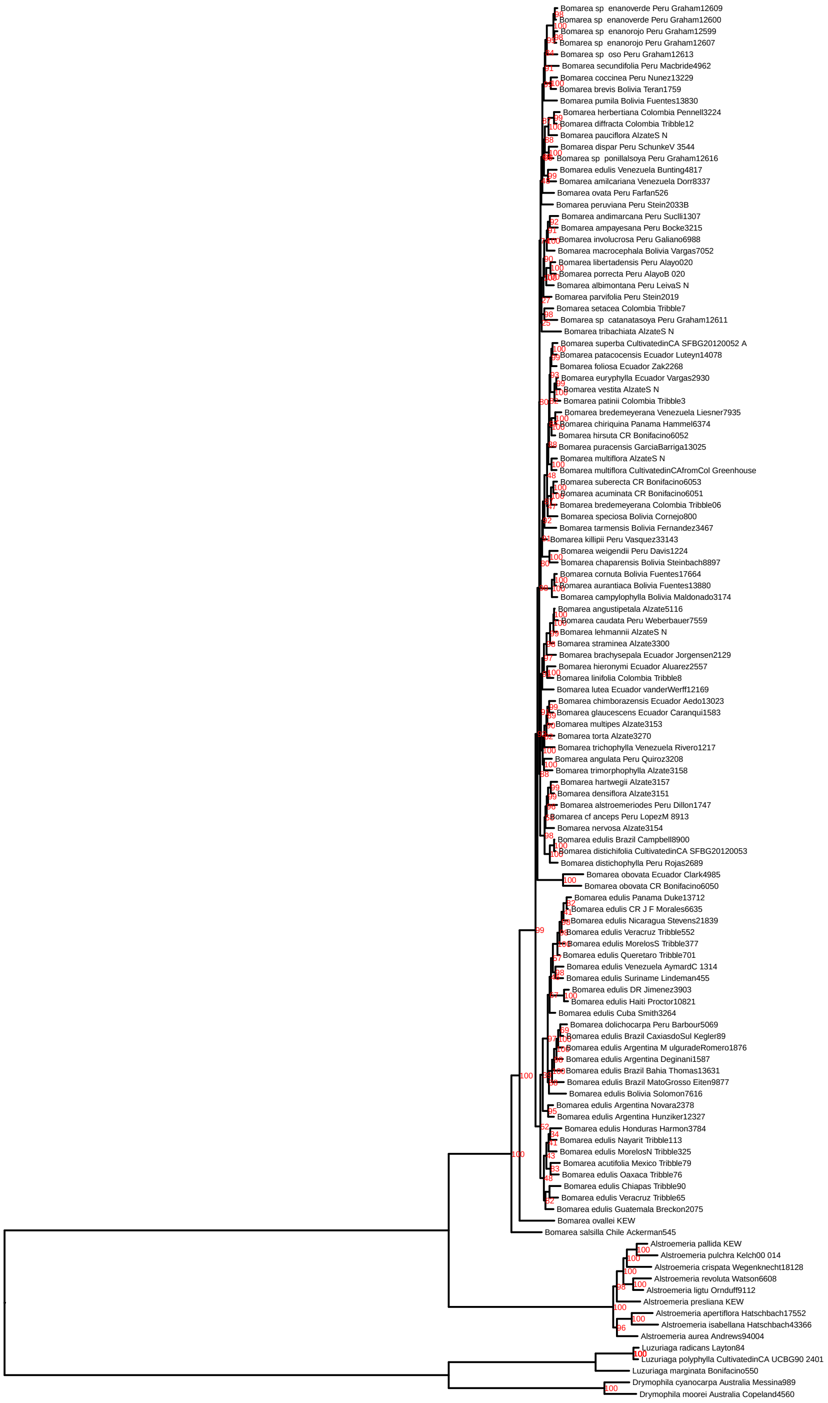


**Figure S2:** Phylogeny produced by the maximum likelihood analysis in `IQtree`
